# High-resolution microscopy of whole plant cells reveals virus-induced changes to intercellular connectivity

**DOI:** 10.64898/2026.09.28.755083

**Authors:** Brandon C. Reagan, Allyson Angermeier, Samantha P. Nuzzi, Mazen Alazem, Andrea A. Zanini, Alejandro Lugo Saaverda, Colten Nichols, Lolita Rotkina, Kirk J. Czymmek, Tessa M. Burch-Smith

## Abstract

Plasmodesmata (PD) provide plant viruses a direct route for cell-to-cell spread. Viral infection increases intercellular trafficking via PD, termed gating, but the mechanism involved remains unclear. Here, using a combination of high-resolution volume electron microscopy and live cell imaging, we demonstrate that diverse viruses increased the number of PD in infected leaves. This increase was triggered by viral movement proteins and the induction of PD formation depended on the presence of the virus and the localization of movement proteins to PD. Further, Group I Remorins, known inhibitors of virus infection, are negative regulators of PD formation and inhibit virus or movement protein-induced changes to PD density. Our results lead to a model in which viral gating of PD may result from increased *de novo* PD formation.

## Main Text

Plant viruses are a persistent threat to global food security, with tens of billions of dollars in annual global crop losses. Understanding how viruses disseminate within their plant hosts is therefore imperative for developing crops with resistance. Unlike animal viruses, plant viruses do not spread by budding from the surface of infected cells but instead move between cells via plasmodesmata (PD), membrane-lined cell-wall-spanning structures that constitute direct cytoplasmic connections between neighboring cells (*1-4*). Despite the long-studied relationship between viruses and PD, relatively little is known about the mechanistic details of how viruses alter PD and their functions. Most plant viruses encode specialized proteins called movement proteins (MPs) that mediate trafficking of the viral genome to and through the PD as ribonucleoprotein complexes or intact virions (*4, 5*). The MPs also mediate a process referred to as gating, where PD in virus-infected cells have increased trafficking capacity compared to their uninfected counterparts. Gating was first characterized for the Tobacco mosaic virus (TMV) 30-kDa movement protein (hereafter MP30) (*6*). PD gating is dynamic, occurring at the leading edge of infection with trafficking reverting to normal away from the infection front (*7*). The leading model for gating invokes degradation of callose, a cell wall polysaccharide, at PD leading to dilation of the pores to increase intercellular flux (*8*). Notably, high-resolution imaging by electron microscopy fails to reveal dilated PD pores in plants expressing MP30 (*9*).

Further, recent evidence suggests that PD size is not a reliable indicator of trafficking capacity as PD with small cytoplasmic volumes may exhibit increased trafficking compared to PD with larger cytoplasmic volumes (*10*). Collectively, these findings suggest that processes besides PD dilation are likely involved in gating.

Leveraging state-of-the-art volume electron microscopy (vEM) and live cell imaging, we report that diverse viruses increase the number of PD and alter their distribution in cells of infected leaves, updating previous findings (*11*). This induction is dependent on the presence of the virus and is not likely a consequence of non-cell-autonomous signaling. Viral MPs alone are sufficient to alter pore formation in the absence of infection, and PD localization is required to induce PD formation and increased trafficking capacity. Transient knockdown of *Nicotiana benthamiana* genes encoding Group I Remorins, previously identified as negative regulators of viral PD gating (*12-14*), revealed that NbREM1 is a negative regulator of PD formation. These findings lead to a new model of PD gating through the altered formation of PD and propose the existence of endogenous mechanisms that regulate PD formation and distribution in cell walls.

## Results

### Tobacco mosaic virus infection results in increased numbers of PD and clustering

PD are integral components of the plant cell wall and cannot be studied apart from this environment. vEM was therefore adopted for *in situ* investigation of cell walls and PD in leaves of *N. benthamiana* plants infected with TMV. Tissue samples from uninfected or systemically infected leaves were prepared for plasma focused ion beam-scanning electron microscopy (pFIB-SEM) (*15*). Iterative milling and imaging (Materials and Methods, fig. S1) were used to capture volumes of whole cells or large portions of whole cells (Fig. 1, fig. S2, movies S1and S2). For the uninfected cells shown in Fig 1, voxels were 7 nm x 7 nm x 10 nm and the data set comprised approximately 2,300 images, representing a volume of more than 8,200 μm^3^. For the virus-infected cells in Fig. 1, a similar number of images with voxel sizes of 8 nm x 8 nm x 10 nm were collected for a volume of almost 16,000 μm^3^ (Fig. 1, fig S2, and movie S2). In the reconstructed volumes produced from these high-resolution pFIB-SEM data sets, organelles were readily visible and the presence of PD was easily determined in both epidermal-epidermal and epidermal-mesophyll cell walls (Fig. 1c,e,g,h). To increase the throughput of pFIB-SEM, Adaptive Scanning, an artificial intelligence (AI) module (*16*) was used to acquire large cell volumes from virus-infected cells (Fig. 1 and fig. S2). In this reconstruction, vacuolar contents are collected at low resolution while retaining high-resolution imaging of the cell walls and the cytoplasm including the viroplasm in TMV-infected cells that appeared to displace most of the ER (fig. S3). pFIB-SEM with Adaptive Scanning reduced collection times from >100 hours to approximately 70 hours for a whole cell (*16*). PD were readily identified in the cell walls of cells whose volumes were captured by Adaptive Scanning FIB-SEM (Fig. 1h and j, movie S2).

**Fig. 1.**
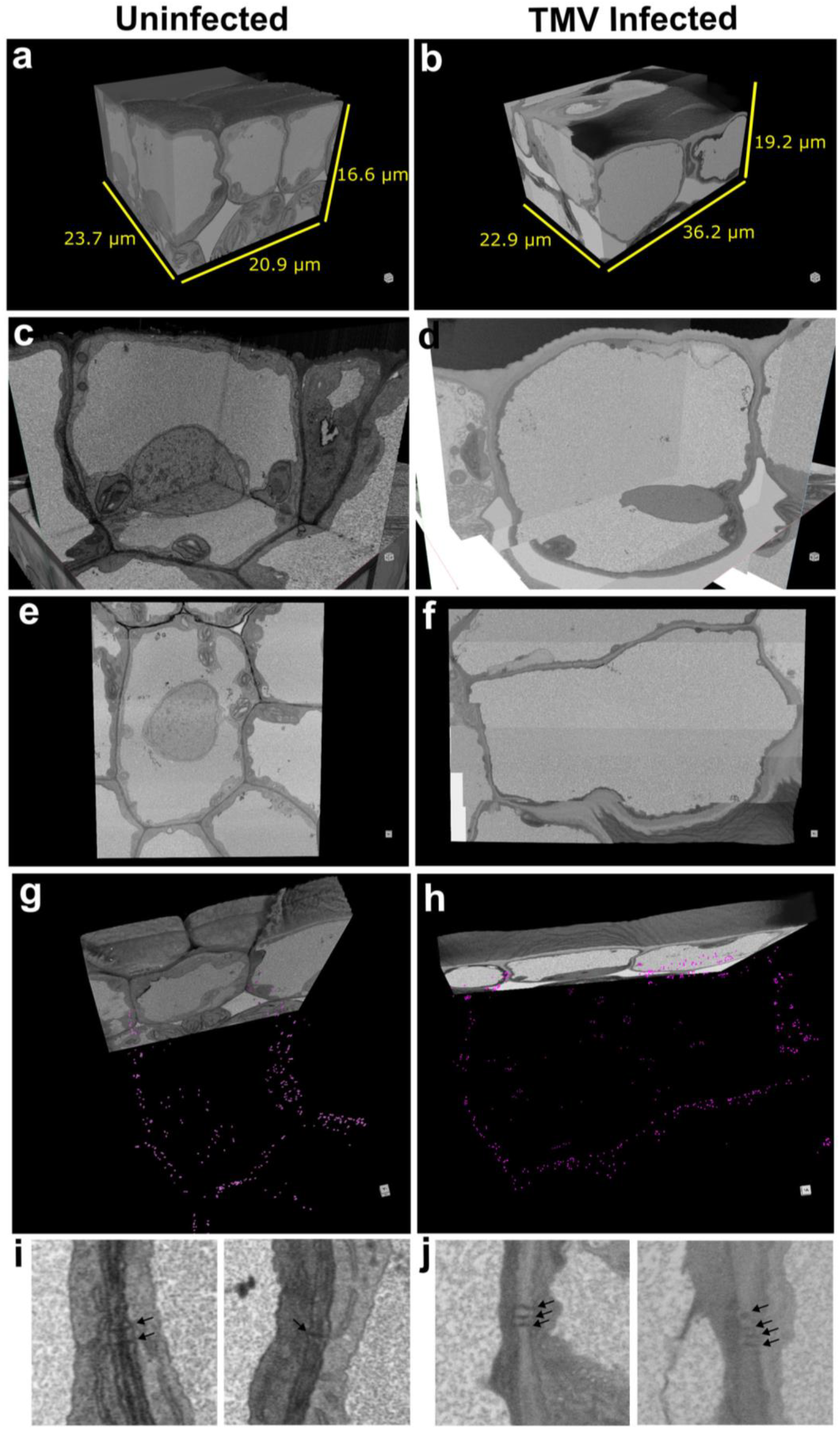
vEM imaging of uninfected and TMV infected tobacco leaf epidermis whole cell. (**A**) 3D view of uninfected and (B) TMV-infected vEM datasets revealing overall epidermal cell structure. (C) 3D orthoslice view of uninfected tissue reveal normal cellular structures, such as the nucleus and chloroplasts. (D) 3D orthoslice view of the TMV-infected tissue reveals a viroplasm indicative of infection (top). (E) A view of the uninfected epidermal cell from the Y-axis revealing the overall whole cell shape. (F) Same as (E) for TMV-infected epidermal cell. (G) 3D view of the uninfected whole epidermal cell with the images partially retracted revealing PD marks (magenta dots). (H) Same as (G) for the TMV-infected epidermal whole cell. (I) Close-up view of two regions of epidermal-epidermal cell wall from volume shown in (A) with arrows pointing to PD. (J) Close-up view of two regions of epidermal-epidermal cell wall in virus-infected cells from volume shown in (B) with arrows pointing to PD.

Using whole cell walls collected by FIB-SEM, individual PD were identified and scored (Fig. 2a-f, colored dots). Points pattern analysis revealed that in both treatments, PD occurred in clusters, where a cluster was defined as a group of three or more PD (Fig. 2g, h and fig. S4). The total number of PD in infected cells was larger than the total number of PD in uninfected cells, resulting in a PD density of ∼0.6 PD μm^-2^ in infected cells compared to ∼0.2 PD μm ^-2^ in uninfected cells (Fig 2i). Consistent with this, the average number of PD in a cluster in infected cells (∼8) was more than double that in uninfected cells (∼3) (Fig. 2j-l) and there were more PD clusters in the infected cells (fig. S4). These results suggest that viral infection modified the cellular machinery that directs the formation of PD in cell walls, leading to larger PD clusters and more PD in a cell wall. In the epidermal-mesophyll cell walls, the organization of PD into clusters was even more striking, with most PD occurring as part of clusters and markedly larger clusters were observed in TMV-infected tissues (fig. S5).

**Fig. 2.**
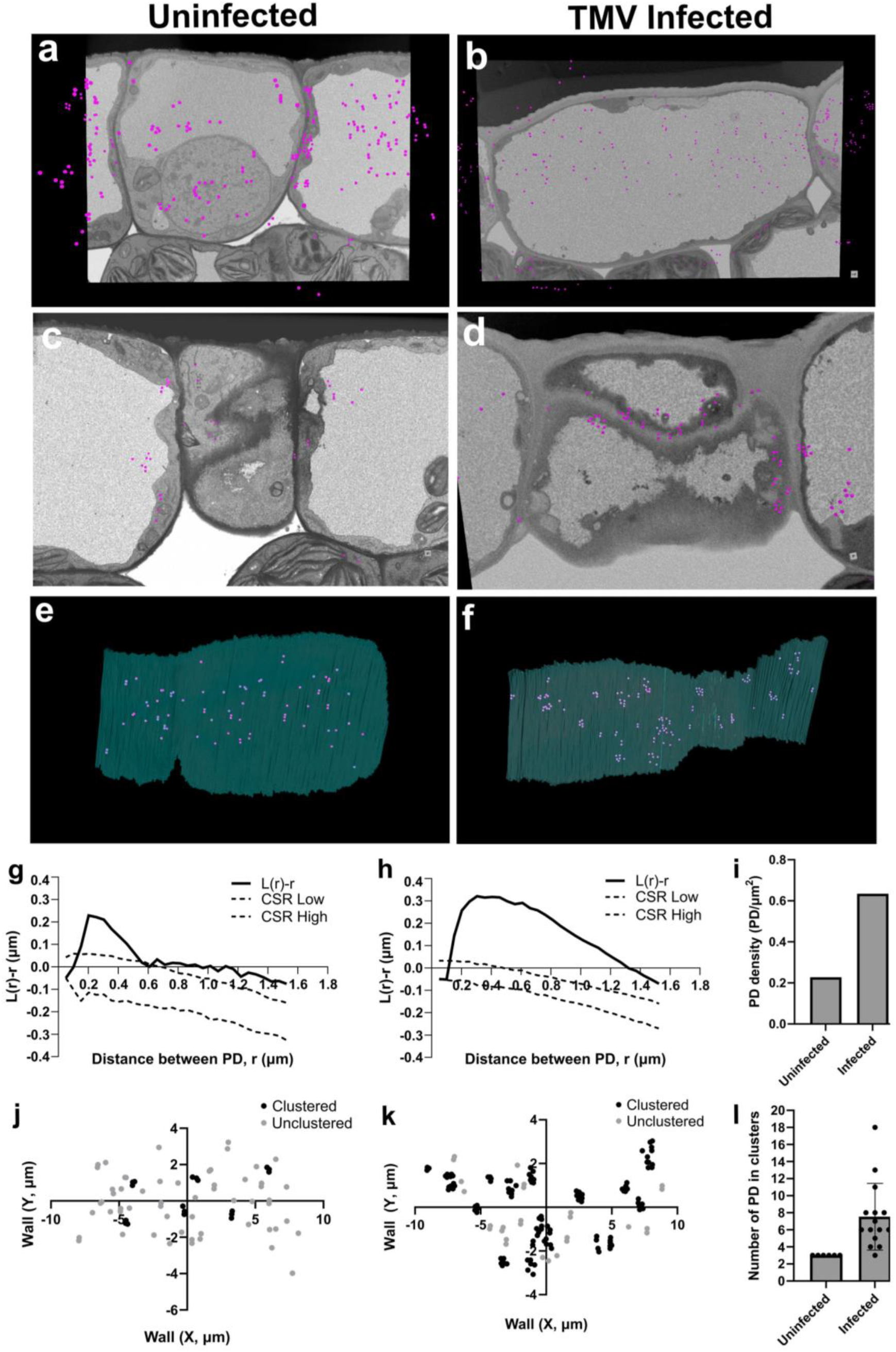
Plasmodesmata distribution in uninfected and TMV-infected epidermal-epidermal cell wall interface. (**A**) vEM images in the Z plane of uninfected and (B) infected epidermal cell overlaid with PD (magenta dots). (C) Merging epidermal cell interfaces of uninfected cells. (D) Same as (C) with TMV-infected epidermal cell. (E) Manual segmentation of an uninfected epidermal-epidermal cell wall interface with PD (magenta dots). (F) Same as (e) for TMV-infected epidermal-epidermal cell wall interface. (G) Spatial analysis of PD distribution on the epidermal-epidermal cell wall interface for uninfected tissue and (H) TMV-infected tissue. Ripley’s L-function is plotted as L(r) – r against the inter-PD distance (r) (solid line in (G) and (H)). The upper and lower bounds of the complete spatial randomness (CSR) region determined by Monte Carlo simulations (95% confidence) are indicated by dashed lines. (I) PD density per unit cell wall area in uninfected and infected epidermal-epidermal cell wall interface. (J) Positions of PD on the uninfected epidermal cell wall interface. PD are classified as “clustered” (black) or “unclustered” (grey) using DBSCAN. (K) Same as (I) for the TMV-infected epidermal cell wall interface with “clustered” in magenta and “unclustered” in grey. (L) Number of PD per cluster in uninfected and TMV-infected epidermal cell walls. Bars represent SD, individual points represent a PD cluster (n=1).

Taking an independent approach to investigating viral effects on PD, the number and distribution of PD in tissues was measured by transmission EM (TEM, (*17*)). Similar to the pFIB-SEM approach, the number of PD connecting neighboring epidermal cells in systemically infected leaves and uninfected *N. benthamiana* leaves was determined. Consistent with the vEM data, TMV infection resulted in an increase in the number of PD in systemically infected leaves with the PD count frequency (F, (*17*)) increasing from 1.5 in uninfected epidermal cells to 3.3 in infected cells (Fig. 3a and b).

**Fig. 3.**
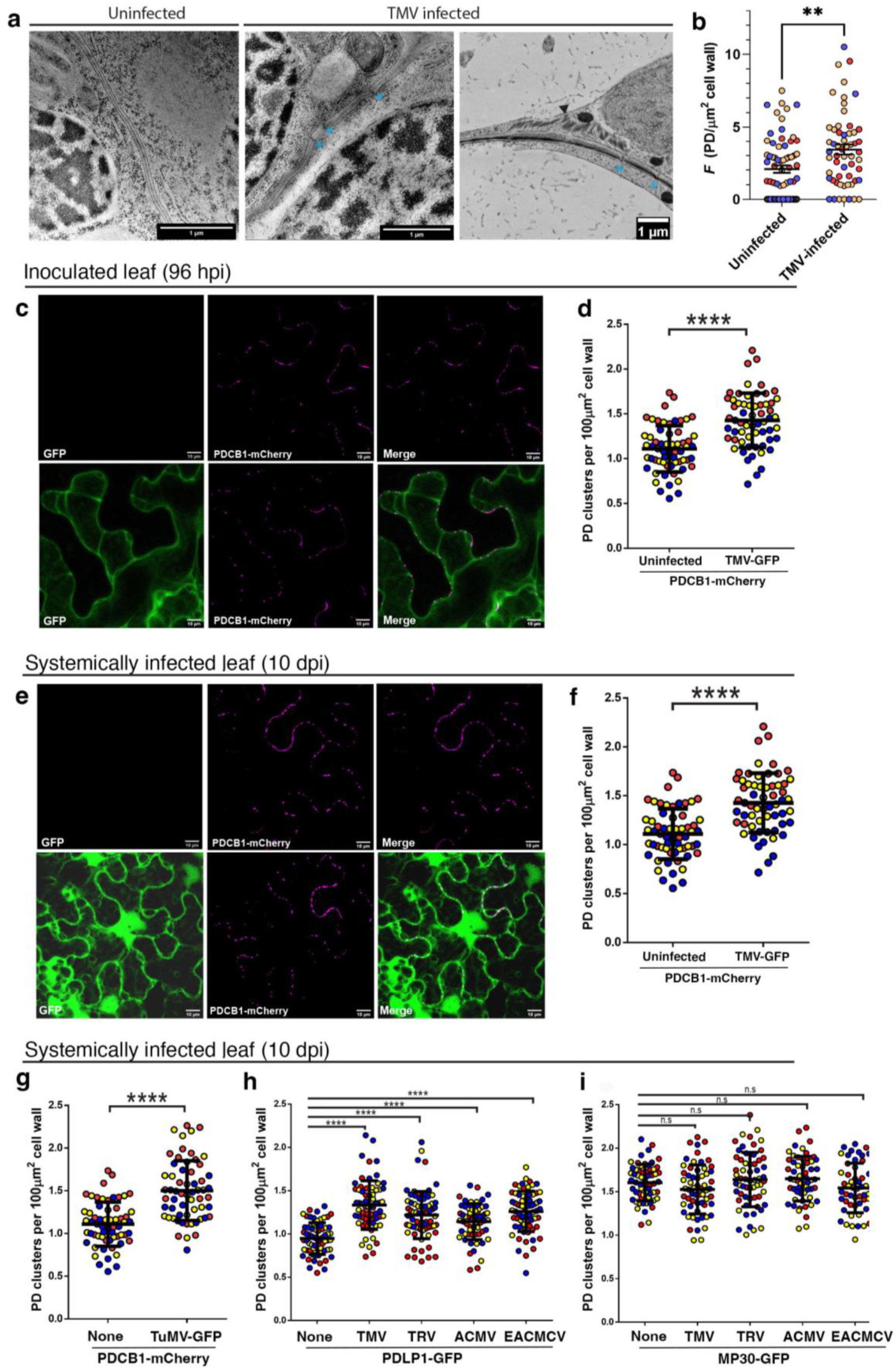
Increased numbers of PD clusters is a conserved feature of viral infection and it depends on the presence of the virus. **(A)** Representative TEM images of uninfected or TMV infected *N. benthamiana* epidermal cells. PD are marked with asterisks; arrowhead points out viroplasm in an infected cell. (B) PD frequency, F the number of PD per μm^2^ of cell wall, was calculated based on TEM imaging of PD. Statistical significance was determined by Mann-Whitney (p < 0.01). Different colors correspond to different biological replicates. (C, E) Representative maximum projection of z-stack of images of *N. benthamiana* epidermal leaf cells expressing PDCB1-mCherry and/or TMV expressing monomeric GFP (TMV-GFP). Scale bar = 10μm. Images were collected from the inoculated leaf 96 hours post infection (hpi) and from systemically infected leaves 10 days post infection (dpi). (D, F) PD cluster densities in inoculated (D) or systemic leaves (F) from uninfected or TMV-GFP infected plants were calculated using AtPDCB1-mCherry as a marker for PD clusters. Cluster densities were quantified for at least 20 fields of view for 3 biological replicates (red, yellow and blue circles represent different biological replicates). (G) PD cluster densities were quantified in plants systemically infected with TuMV-GFP 10 dpi using PDCB1-mCherry as a PD cluster marker. (H) PD cluster densities were quantified in plants systemically infected with TMV, TRV, ACMV, or EACMCV 10 dpi using PDLP1-GFP as the PD marker. (I) Densities of PD clusters in leaves expressing MP30-GFP while uninfected or systemically infected with a virus. MP30-GFP was used as a PD marker. PD cluster densities for 20 foci for each of 3 biological replicates are shown (red, blue, and yellow circles). Statistical significance was determined using Student’s t-test (p < 0.0001).

One drawback of EM is the low sample throughput. PLASMODESMATA LOCATED PROTEIN 1 fused with GFP (AtPDLP1-GFP) and PLASMODESMATA CALLLOSE BINDING1 fused with mCherry (AtPDCB1-mCherry) are established PD markers (*18, 19*), localizing to foci in the cell wall that denote the presence of PD clusters. AtPDLP1-GFP and AtPDCB1-mCherry were adopted for measuring PD density via confocal microscopy, a more rapid and high throughput imaging modality (fig. S6). *N. benthamiana* leaves were co-inoculated with Agrobacterium carrying constructs for expression of AtPDCB1-mCherry and TMV expressing free GFP (TMV-30B) or an empty vector as a control. After 80 hours of transgene expression and virus infection, PD density was measured in epidermal cells of inoculated leaves by scoring AtPDCB1-mCherry foci in the presence and absence of TMV-GFP infection (Fig. 3c-f). The total number of fluorescent foci per unit cell wall length in a 15-μm z-stack was used to calculate PD cluster density (Fig. S6, (*20*)). Consistent with the EM results, TMV-GFP infected epidermal cells exhibited a mean of 1.2 clusters per 100 μm^2^ cell wall compared to 0.8 clusters per 100 μm^2^ cell wall in uninfected control cells, representing a statistically significant increase (43.9%) in the number of fluorescent foci (Fig. 3c and d). A similar increase in the number of AtPDCB1-mCherry foci was observed in systemically infected leaves (Fig. 3e and f). While the epidermal cells of upper leaves of uninfected plants contained 1.1 clusters per 100 μm^2^ cell walls, epidermal cells of virus-infected systemic leaves had an average of 1.4 clusters per 100 μm^2^ cell wall, representing a 28.7% increase in the number of PD (Fig. 1d). This increase was dependent on the presence of the virus in the tissue since there was no increase in fluorescent foci before systemic infection by the virus (fig. S7).

### PD formation is a conserved feature of virus infection and requires the presence of the virus

To determine if induction of PD formation is a conserved strategy used by diverse plant viruses to facilitate their cell-to-cell spread, *N. benthamiana* plants were infected with unrelated viruses that move cell-to-cell as ribonucleoprotein complexes or in vesicles, i.e., without the formation of tubules. For this, *N. benthamiana* plants were infected with Turnip mosaic virus (TuMV-GFP, potyviridae) and Tobacco rattle virus (TRV, Tobraviridae). The ability of MPs from phloem-associated viruses to affect PD outside the phloem was examined by infection with either African cassava mosaic virus (ACMV) or East African cassava mosaic Cameroon virus (EACMCV). The presence of the virus in systemic leaves was confirmed by PCR (Fig. S8). PD distributions in systemic leaves infected with TuMV-GFP were quantified using AtPDCB1-mCherry as for TMV-GFP. An increase from 1.1 clusters per 100 μm^2^ cell wall in uninfected control leaves to 1.5 clusters per 100 μm^2^ cell wall in infected leaves was observed (Fig. 3g). PD densities were quantified using PDLP1-GFP when unlabeled viruses were tested. Consistent with our findings with TMV-GFP and TuMV-GFP, systemically infected leaves with every virus tested had elevated numbers of PD clusters compared to uninfected controls: a density of 0.95 clusters per 100 μm^2^ cell wall in the uninfected controls compared to 1.3, 1.2, 1.1 and 1.25 clusters per 100 μm^2^ cell wall for TMV, TRV, ACMV and EACMCV infections, respectively. Thus, diverse families of viruses induce PD formation (Fig. 3g, h). Notably, expression of MP30-GFP in leaves systemically infected with different viruses resulted in no significant change in PD density compared to uninfected controls (Fig. 3i), revealing no synergism or enhancement of the effects of the native viral MPs by MP30.

### The TMV movement protein induces PD formation

Viral MPs gate PD during infection, facilitating viral-cell-to-cell movement and leading to increased intercellular trafficking of unrelated molecules (*4, 5*). To determine if the increase in cluster densities observed in TMV-GFP infected leaves was initiated by the TMV MP, MP30 fused to GFP (MP30-GFP) was co-expressed with AtPDCB1-mCherry in virus-free *N. benthamiana* leaves. The PD density in the presence of MP30-GFP was 1.1 clusters per 100 μm^2^ of cell wall compared to 0.8 in leaves without MP30-GFP, a statically significant increase of 28.4% (Fig. 4a-c). Given that MP30 localizes to PD and has been used extensively as a marker for PD (*4*), PD cluster densities were measured in epidermal cells expressing MP30-GFP alone (Fig. 4d). The MP30-cluster density of 1.1 cluster per 100 μm^-2^ of cell walls was not statistically significantly different from the density observed with the AtPDCB1-mCherry marker in the presence of MP30 but represented a significant increase over cluster density in cells without MP30 (Fig. 4c). Thus, TMV MP30 alone is sufficient to induce PD formation. Potato leafroll virus (PLRV) is restricted to the phloem during infections of its plant hosts. Nevertheless, its MP, MP17, can gate PD of leaf epidermal cells (*21*). Like MP30, MP17 alone was able to induce PD formation in *N. benthamiana* leaf epidermal cells (Fig. S9). Induction of *de novo* PD formation may be a general property of MPs.

**Fig. 4.**
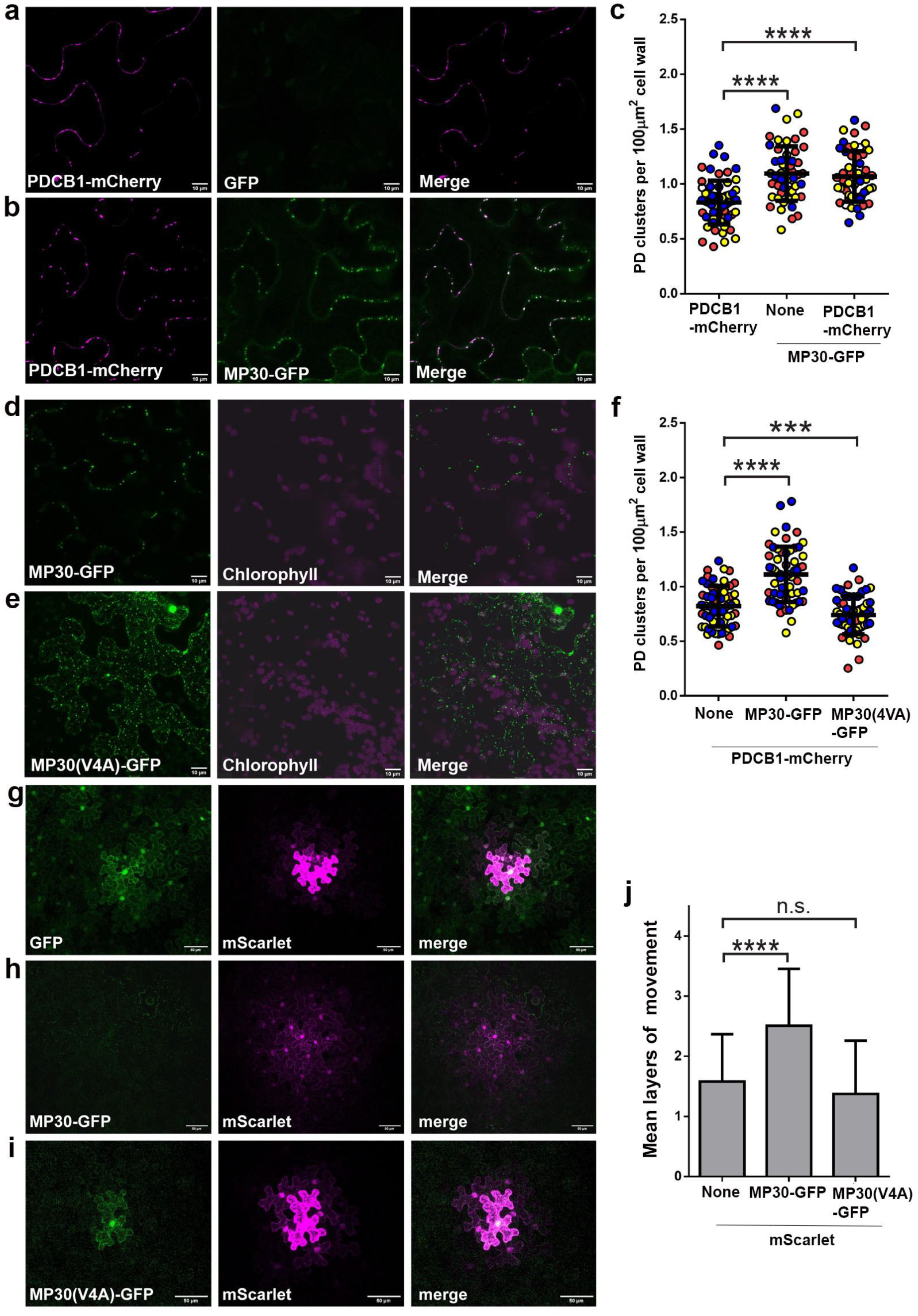
The TMV movement protein induces PD formation. (**A**) Representative maximum projection of *z*-stack of images of epidermal cells expression either PDCB1-mChery alone or (B) with MP30-GFP are shown. Scale bars = 10μm. (C) PD cluster densities were calculated for both PDCB1-mChery and MP30-GFP expressed alone or together. (D,E) Representative images of *N. benthamiana* leaf epidermal cells expressing WT MP30-GFP (D) or the mutant MP30(V4A)-GFP (E). Scale bars = 10μm. (F) PD cluster densities in the presence of MP30-GFP or MP30(V4A)-GFP were determined using PDCB1-mCherry as the PD marker were calculated for at least 20 fields of view for 3 biological replicates (red, blue, and yellow circles). Statistical significance was determined by Student’s t-test (p < 0.001). (G-I) Representative maximum projection of *z*-stack of images of epidermal cells expressing soluble mScarlet in the presence of GFP, MP30-GFP, MP30(V4A) that were used to measure intercellular trafficking. (J) The mean layers of movement (intercellular trafficking) of mScarlet plus the standard deviation for foci sored in (G-I) were calculated. Statistical significance was determined using the Mann-Whitney U-test or bootstrap method (p < 0.001).

To determine if PD localization of MP30 is required for the increase in PD density, a mutation previously shown to block PD localization of MP30 was introduced into MP30-GFP to yield MP30(V4A)-GFP (*22*). In *N. benthamiana* leaf epidermal cells, the MP30(V4A)-GFP variant failed to localize to PD and instead accumulated in other organelles (Fig. 4e). MP30(V4A)-GFP did not induce PD formation and instead there was a slight decrease in AtPDBC1-mCherry-labeled clusters (Fig. 4f). Further, while wild-type MP30 enhanced the trafficking of soluble mScarlet in a movement assay, MP30(V4A)-GFP had no effect on intercellular trafficking of mScarlet (Fig. 4g-j). We interpret this to mean that the mutant MP is unable to gate PD and increase intercellular trafficking and is also unable to induce PD formation. Thus, PD localization of MP30 is required to both induce PD formation and to gate PD.

### NbREM1 is a negative regulator of PD formation

Group I Remorins (REMs) are negative regulators of virus-induced gating of PD (*13, 20*) (*12, 14*). To investigate the role of Group 1 NbREM1 in inhibiting gating, an RNA silencing construct that targeted both *N. benthamiana* homologs, *NbREM1*.*1* and *NbREM1*.*2*, was used to simultaneously knockdown their expression by virus-induced gene silencing (VIGS, fig. S10). Consistent with previous work with *rem* mutant plants (*23*), *NbREM1*-silenced plants exhibited increased intercellular trafficking of soluble GFP compared to non-silencing control plants (fig. S11). Expression of PDLP1-GFP in silenced plants revealed that these plants had elevated numbers of PD clusters (1.3 per 100 μm^2^ cell wall) compared to non-silencing controls (1.1 per 100 μm^2^ cell wall, Fig. 5a and b). Next, the effects of reduced *NbREM1* expression on MP-mediated PD formation were measured. The number of MP30-labeled clusters in *NbREM1*-silenced plants was increased compared to non-silenced control plants (1.6 vs 1.2 clusters per 100 μm^2^ cell wall, Fig. 5c and b). This suggests that NbREM1 also restricts MP-induced PD formation.

**Fig. 5.**
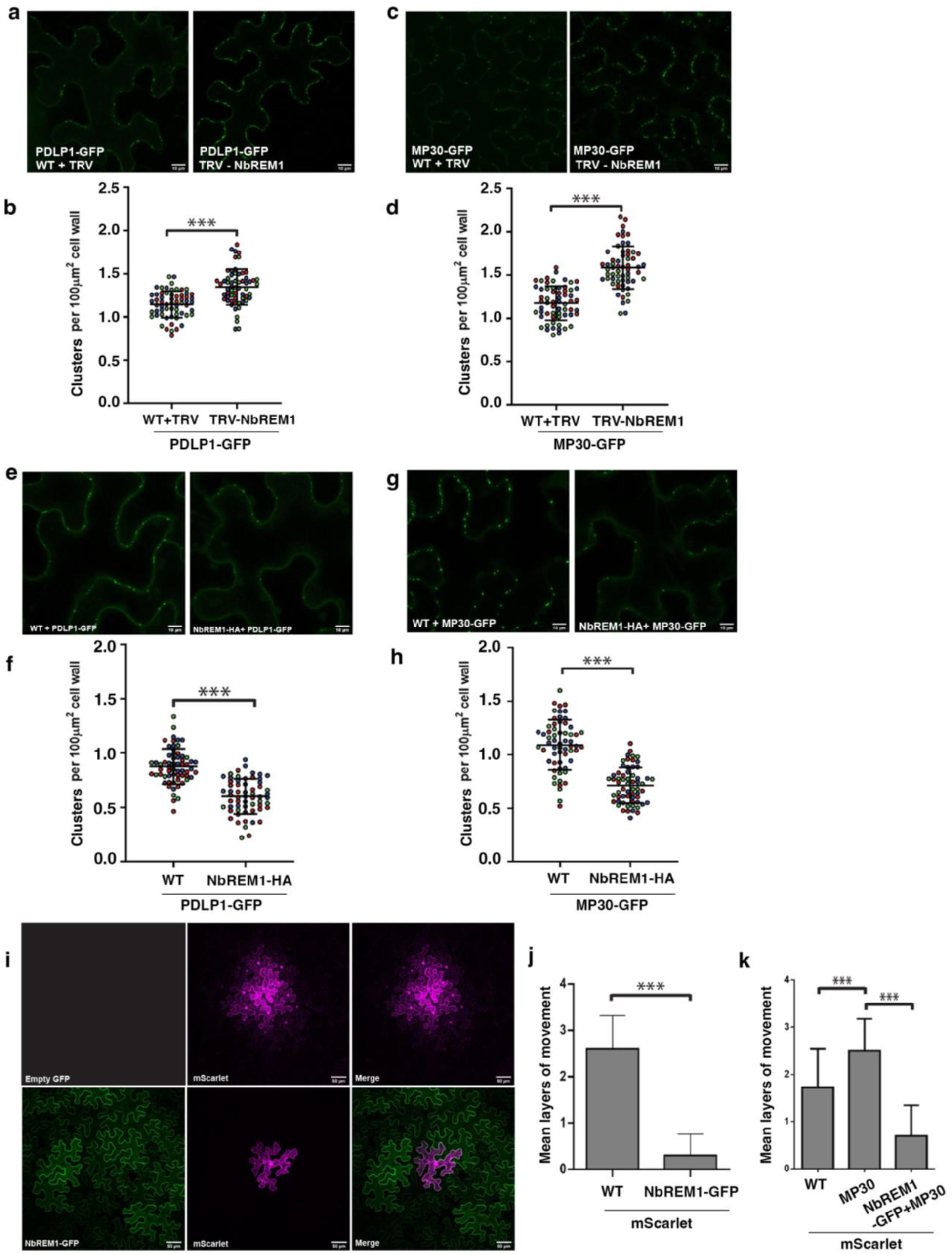
NbREM1 is a negative regulator of PD formation. (**A**) Representative maximum projection of *z*-stack of images of cells expressing PDLP1-GFP in non-silencing controls (left) or in *NbREM1*-silenced plants (right) are shown. Scale bars = 10 μm. (B) PD cluster densities for 20 fields of view for each of 3 biological replicates (red, blue, and green circles) are shown for PDLP1-GFP. Student’s t-test (p < 0.001). (C) Representative maximum projection of *z*-stack of images of cells expressing MP30-GFP in non-silencing controls (left) or in *NbREM1*-silenced plants (right). Scale bars = 10 μm. (D) PD cluster densities for 20 fields of view for each of 3 biological r replicates (red, blue, and yellow circles) are shown. Statistical significance was determined using a Student’s t-test (p < 0.001). (E, F) Representative maximum projection of *z*-stack of images of cluster foci labeled with PDLP1-GFP in WT controls (left in E) and plants overexpressing NbREM1.1-HA (right in E) or labeled with MP30-GFP in WT controls (left in F) and plants overexpressing NbREM1.1-HA (right in F). Scale bars = 10μm. (G, H) PD cluster densities measured with PGLP1-GFP (G) or MP30-GFP (H) for 20 foci for each of three biological replicates. Data from each replicate are shown (red, blue, and green circles). Statistical significance was determined with Student’s t-test (p < 0.001). (I) Intercellular trafficking was measured by counting the number of layers of mScarlet spread in *N. benthamiana* leaf epidermal cells. Representative images of foci in the absence (top row) or presence of NbREM1.1-GFP (bottom row) are shown. Scale bars = 50 μm. (J) Mean layers of movement (intercellular trafficking) and standard deviation were calculated. Data for three biological replicates are shown and at least 20 foci were collected for each biological replicate. Statistical significance was determined using the Mann-Whitney U-test or bootstrap method (p < 0.001).

Overexpression of NbREM1.1 tagged at its C-terminus with the HA epitope tag (NbREM1-HA) from the constitutive 35S promoter (Fig. S10) resulted in decreased numbers of PDLP1-GFP-labeled clusters (0.6 per 100 μm^2^ cell wall) compared to plants with wild-type levels of REM1 (0.9 per 100μm^2^ cell wall, Fig. 5 e and f). Overexpression of NbREM1-HA also resulted in decreased PD densities in plants expressing MP30-GFP (0.7 clusters per 100 μm^2^ vs 1.1 per 100 μm^2^ cell wall in control plants), indicating that overexpression of NbREM1 inhibits MP-induced PD formation (Figure 5g and h).

When similarly overexpressed in *N. benthamiana* leaves, NbREM1.1tagged with GFP (NbREM1-GFP) labeled the plasma membrane as previously reported (Fig. 5i) (*24*). In contrast to *NbREM1*-silenced plants, plants transiently over-expressing NbREM1.1-GFP inhibited intercellular trafficking of the soluble mScarlet probe (Fig. 5j). Further, in leaves co-expressing NbREM1-HA, MP30-GFP, and mScarlet (Fig. 5c and d), MP30 was unable to increase trafficking of mScarlet compared to plants co-expressing only MP30 and mScarlet (Fig. 5k). This result supports the idea that the *de novo* PD induced by MP30 are responsible for the increased trafficking capacity observed in tissues containing MP30, and likely TMV.

## Discussion

Taken together, our data reveal that the increase in trafficking observed during infection is at least in part due to increased numbers of PD, arranged into large clusters, in the infected leaves. This is consistent with evidence supporting a positive correlation between the number of PD in a tissue and its trafficking capacity (*17, 20*). Primary PD formation, where PD form through ER-dependent incomplete cytokinesis, is well documented (*25-28*). In contrast, secondary PD formation in the absence of cytokinesis, remains poorly understood despite being common during cell expansion and development (*29-31*). The PD formed during viral infections are likely secondary PD since we observed changes in PD density and distribution in leaves that were in the expansion phase of their growth (inoculated leaves in Fig. 3 and upper in leaves in Figs. 4 and 5). These virus-induced PD are likely required for the cell-to-cell spread of viral ribonucleoprotein complexes or vesicles, since failure to induce PD resulted in reduced intercellular trafficking (Fig. 4), consistent with numerous previous reports of the requirement of movement proteins for viral infection (*27, 32, 33*). Further experiments will be required to determine whether these PD are distinct from PD formed in the normal course of plant development.

We propose that induction of PD formation is likely a conserved feature of infection by diverse families of plant viruses, as revealed in this work where PD numbers in the presence of potyviruses which do not encode distinct movement proteins, geminiviruses which are expected to be restricted to the vasculature, and other viruses were found to increase PD density during infection (Fig. 1-3). Our data also demonstrate that the TMV movement protein, MP30, is sufficient to induce secondary PD formation to levels similar to those seen in plants infected with TMV (Fig. 4). The MP17 of Potato leafroll virus (a polerovirus) shares general features with MP30 but is distinct from the tobamovirus protein (*34*). MP17 was also able to induce PD formation, (fig. S9), suggesting that PD formation may be a common property of MPs that gate PD.

Our results suggest viruses co-opt existing cellular mechanisms to induce the formation of PD during infection. REMs are plant specific proteins and nanodomain markers and are separated into six groups with certain members having roles in regulating PD function and virus infection (*13*) (*35*); (*36*). Group 1 REMs negatively regulate virus infection. In contrast, Group 4 REMs positively regulate geminivirus infection (*37*). Our data support a role for these proteins in limiting *de novo* pore insertion in both the presence and absence of viral infection. They also point to NbREM1 limiting virus infection by restricting PD formation. Thus, we propose REMs as part of the cellular machinery involved in *de novo* PD formation. Interestingly, NbREM1 is targeted for degradation by the Tomato mosaic virus (ToMV) movement protein (*38*).We posit that the removal of NbREM1 in this context is to permit *de novo* PD formation and support viral cell-to-cell movement. Together, these results point to viruses evolving strategies to circumvent host regulatory networks for determining PD densities.

Introduction of a single amino acid substitution into the plasmodesmata localization signal (PLS) of MP30 (MP30(V4A)), a mutation previously shown to disrupt its localization, prevented the induction of PD formation (Fig. 4). Thus, proper localization of the MP to PD is required to induce PD formation and suggests that the cellular machinery for *de novo* PD formation may be present in the vicinity of existing PD and that the local environment of the cell wall at PD is ideal for pore insertion. This is consistent with the multiple twinning model of secondary PD formation where new PD are formed in proximity to existing PD (*39*). Next, our data reveal that REMs suppress PD formation. REMs localize to and help regulate the organization of PM nanodomains (*36*) and these dynamics can be altered by viral infection (*23, 40, 41*).

In the context of REMs’ roles in plasma membrane nanodomain lipid sequestration and our present findings, our data suggest a new model for viral gating of PD (fig. S12). We propose that specialized organization of the PM is critical in determining sites of *de novo* PD formation and that loss of REMs, caused by the virus or viral movement protein, allows membrane disorder and unrestricted distribution of positive regulators of PD formation. This new distribution of positive regulators of PD formation is permissive for forming increased numbers of PD near existing PD and produced large clusters of PD containing increased numbers of PD (fig. S12). Existing models for how viruses gate PD have largely invoked a role for callose metabolism where removal of PD-associated callose increases the size of the pore to facilitate viral intercellular movement. Our findings do not rule out the involvement of callose in regulating trafficking but instead suggest that there is additional complexity to the intercellular trafficking of viruses that requires currently undescribed cell wall dynamics. Virus-infected tissues will therefore be a crucial model for studying the intriguing process of secondary plasmodesmata formation.

Together, our findings suggest that while the mechanisms used by various viruses to induce PD formation may differ, the induction of PD formation is a conserved step in the spread of virus infection. Potyviruses are the largest family of plant viruses, followed by the geminiviruses. Together these two groups represent some of the most commercially important plant viruses, responsible for staggering agricultural losses each year (*42*). Understanding mechanisms of infection could therefore provide valuable insight into developing new mitigation strategies.

## Acknowledgments

We thank Dr. Vincent Fondong for the infectious clones for ACMV and EACMCV, and Dr. Christine Faulkner for the AtPDCB1-mCherry construct. We also thank Jaydeep Kolape of the UTK Advanced Microscopy Imaging Facility for help with TEM image collection. Members of the Materials and Structural Analysis Division of Thermo Fisher Scientific contributed significantly to the use of the Adaptative Scanning technology used in this study.

## Funding

This work was supported by the National Science Foundation grant MCB1846245 (TBS) and internal funds form the Donald Danforth Plant Science Center.

## Author contributions

Conceptualization: BCR, TBS

Methodology: BCR, AA, LR, KJC, TMBS

Investigation: BCR, AA, AAZ, SPN, MA ALS,

Visualization: BCR, AA, AAZ, ALS

Funding acquisition: TMBS Supervision: KJC, TMBS

Writing – original draft: BCR

Writing – review & editing: BCR, AA, KJC, TMBS

## Competing interests

Authors declare that they have no competing interests.

## Data, code, and materials availability

pFIB-SEM datasets have been deposited in the Electron Microscopy Public Image Archive (EMPIAR) under the accession number EMPIAR-13966. Code for PD analysis has been deposited at GITHUB and is available through Zenodo: 10.5281/zenodo.22812106. All other data are available in the main text or the supplementary materials.

## Supplementary Materials

Materials and Methods

Figs. S1 to S11

Tables S1 and S2

Movies S1 and S2

## Notes

doi:10.5281/zenodo.22812106

### Competing Interest Statement

The authors have declared no competing interest.

### Summary of Updates

Legend to Figure 1 revised and author name corrected

## References

1. E. M. Bayer, Y. Benitez-Alfonso, Plasmodesmata: Channels Under Pressure. Annu Rev Plant Biol 75, 291–317 (2024).

2. E. E. Tee, C. Faulkner, Plasmodesmata and intercellular molecular traffic control. The New phytologist 243, 32–47 (2024).

3. A. A. Zanini, T. M. Burch-Smith, New insights into plasmodesmata: complex ‘protoplasmic connecting threads’. J Exp Bot 75, 5557–5567 (2024).

4. B. C. Reagan, T. M. Burch-Smith, Viruses Reveal the Secrets of Plasmodesmal Cell Biology. Mol Plant Microbe Interact 33, 26–39 (2020).

5. W. J. Lucas, Plant viral movement proteins: agents for cell-to-cell trafficking of viral genomes. Virology 344, 169–184 (2006).

6. S. Wolf, C. M. Deom, R. N. Beachy, W. J. Lucas, Movement protein of tobacco mosaic virus modifies plasmodesmatal size exclusion limit. Science 246, 377–379 (1989).

7. K. J. Oparka, D. A. Prior, S. Santa Cruz, H. S. Padgett, R. N. Beachy, Gating of epidermal plasmodesmata is restricted to the leading edge of expanding infection sites of tobacco mosaic virus (TMV). Plant J 12, 781–789 (1997).

8. B. L. Epel, Plant viruses spread by diffusion on ER-associated movement-protein-rafts through plasmodesmata gated by viral induced host beta-1,3-glucanases. Semin Cell Dev Biol 20, 1074–1081 (2009).

9. P. J. Moore, C. A. Fenczik, C. M. Deom, R. N. Beachy, Developmental changes in plasmodesmata in transgenic tobacco expressing the movement protein of tobacco mosaic virus. Protoplasma 170, 115–127 (1992).

10. D. Yan et al., Sphingolipid biosynthesis modulates plasmodesmal ultrastructure and phloem unloading. Nat Plants 5, 604–615 (2019).

11. B. C. Reagan, S. P. Nuzzi, A. A. Zanini, T. M. Burch-Smith, Plant Viruses Induce Plasmodesmata Formation During Infection. SSRN, (2023).

12. S. Fu et al., Rice Stripe Virus Interferes with S-acylation of Remorin and Induces Its Autophagic Degradation to Facilitate Virus Infection. Mol Plant 11, 269–287 (2018).

13. S. Raffaele et al., Remorin, a solanaceae protein resident in membrane rafts and plasmodesmata, impairs potato virus X movement. Plant Cell 21, 1541–1555 (2009).

14. G. Cheng, Z. Yang, H. Zhang, J. Zhang, J. Xu, Remorin interacting with PCaP1 impairs Turnip mosaic virus intercellular movement but is antagonised by VPg. The New phytologist 225, 2122–2139 (2020).

15. J. S. Wickramanayake, K. J. Czymmek, in Methods in Cell Biology, K. Narayan, L. Collinson, P. Verkade, Eds. (Academic Press, 2023), vol. 177, pp. 83–99.

16. T. Hurník Konečná et al., Contextual High-Throughput 3D Volume Electron Microscopy Data Acquisition Using Artificial Intelligence. Microscopy and microanalysis : the official journal of Microscopy Society of America, Microbeam Analysis Society, Microscopical Society of Canada 32, (2026).

17. T. M. Burch-Smith, P. C. Zambryski, Loss of INCREASED SIZE EXCLUSION LIMIT (ISE)1 or ISE2 increases the formation of secondary plasmodesmata. Curr Biol 20, 989–993 (2010).

18. J. Fitzgibbon et al., A developmental framework for complex plasmodesmata formation revealed by large-scale imaging of the Arabidopsis leaf epidermis. Plant Cell 25, 57–70 (2013).

19. C. Simpson, C. Thomas, K. Findlay, E. Bayer, A. J. Maule, An Arabidopsis GPI-anchor plasmodesmal neck protein with callose binding activity and potential to regulate cell-to-cell trafficking. Plant Cell 21, 581–594 (2009).

20. E. E. Ganusova et al., Chloroplast-to-nucleus retrograde signalling controls intercellular trafficking via plasmodesmata formation. Philos Trans R Soc Lond B Biol Sci 375, 20190408 (2020).

21. D. Hofius et al., Evidence for expression level-dependent modulation of carbohydrate status and viral resistance by the potato leafroll virus movement protein in transgenic tobacco plants. The Plant Journal 28, 529–543 (2001).

22. C. Yuan, S. G. Lazarowitz, V. Citovsky, Identification of a Functional Plasmodesmal Localization Signal in a Plant Viral Cell-To-Cell-Movement Protein. MBio 7, e02052–02015 (2016).

23. D. Huang et al., Salicylic acid-mediated plasmodesmal closure via Remorin-dependent lipid organization. Proc Natl Acad Sci U S A 116, 21274–21284 (2019).

24. C. L. Thomas, E. M. Bayer, C. Ritzenthaler, L. Fernandez-Calvino, A. J. Maule, Specific targeting of a plasmodesmal protein affecting cell-to-cell communication. PLoS Biol 6, e7 (2008).

25. L. Wegner, C. Herrfurth, I. Feussner, K. Ehlers, T. M. Haslam, Complex sphingolipid metabolism impacts cell division and plasmodesmal development in the moss Physcomitrium patens. Plant Physiol 199, (2025).

26. Z. P. Li et al., Plant plasmodesmata bridges form through ER-dependent incomplete cytokinesis. Science 386, 538–545 (2024).

27. Y. Kan, V. Citovsky, The roles of movement and coat proteins in the transport of tobamoviruses between plant cells. Frontiers in plant science 16, 1580554 (2025).

28. M. L. Brault et al., Multiple C2 domains and transmembrane region proteins (MCTPs) tether membranes at plasmodesmata. EMBO Rep 20, e47182 (2019).

29. K. Ehlers, R. Kollmann, Primary and secondary plasmodesmata: structure, origin, and functioning. Protoplasma 216, 1–30 (2001).

30. T. M. Burch-Smith, S. Stonebloom, M. Xu, P. C. Zambryski, Plasmodesmata during development: re-examination of the importance of primary, secondary, and branched plasmodesmata structure versus function. Protoplasma 248, 61–74 (2011).

31. K. Aoki, A. Tsushima, Interspecific secondary plasmodesmata at the parasitic interface. Plant Cell Physiol 67, 460–468 (2026).

32. M. Alazem, S. N. Nuzzi, T. M. Burch-Smith, Regulation of cell-to-cell trafficking by viral movement proteins. Journal of experimental botany, (2025).

33. A. M. Wang, Cell-to-cell movement of plant viruses via plasmodesmata: a current perspective on potyviruses. Current Opinion in Virology 48, 10–16 (2021).

34. S. L. DeBlasio et al., The Interaction Dynamics of Two Potato Leafroll Virus Movement Proteins Affects Their Localization to the Outer Membranes of Mitochondria and Plastids. Viruses 10, (2018).

35. I. K. Jarsch, T. Ott, Perspectives on remorin proteins, membrane rafts, and their role during plant-microbe interactions. Mol Plant Microbe Interact 24, 7–12 (2011).

36. P. Gouguet et al., Connecting the dots: from nanodomains to physiological functions of REMORINs. Plant Physiol 185, 632–649 (2021).

37. S. T. Lilly, R. S. Drummond, M. N. Pearson, R. M. MacDiarmid, Identification and validation of reference genes for normalization of transcripts from virus-infected Arabidopsis thaliana. Mol Plant Microbe Interact 24, 294–304 (2011).

38. N. Sasaki, E. Takashima, H. Nyunoya, Altered Subcellular Localization of a Tobacco Membrane Raft-Associated Remorin Protein by Tobamovirus Infection and Transient Expression of Viral Replication and Movement Proteins. Frontiers in plant science 9, 619 (2018).

39. C. Faulkner, O. E. Akman, K. Bell, C. Jeffree, K. Oparka, Peeking into pit fields: a multiple twinning model of secondary plasmodesmata formation in tobacco. Plant Cell 20, 1504–1518 (2008).

40. A. Perraki et al., REM1.3’s phospho-status defines its plasma membrane nanodomain organization and activity in restricting PVX cell-to-cell movement. PLoS pathogens 14, e1007378 (2018).

41. M. D. Jolivet et al., Interdependence of plasma membrane nanoscale dynamics of a kinase and its cognate substrate underlies Arabidopsis response to viral infection. eLife 12, (2025).

42. S. K. Sastry, T. A. Zitter, in Plant Virus and Viroid Diseases in the Tropics Vol. 2 Epidemiology and Management (Springer, Dordrecht, 2014), vol. 2: Epidemiolgy and Management, pp. 149–480.

43. Y. Hua, P. Laserstein, M. Helmstaedter, Large-volume en-bloc staining for electron microscopy-based connectomics. Nature communications 6, 7923 (2015).

44. C. A. Schneider, W. S. Rasband, K. W. Eliceiri, NIH Image to ImageJ: 25 years of image analysis. Nature methods 9, 671–675 (2012).

45. Y. Liu, T. Burch-Smith, M. Schiff, S. Feng, S. P. Dinesh-Kumar, Molecular chaperone Hsp90 associates with resistance protein N and its signaling proteins SGT1 and Rar1 to modulate an innate immune response in plants. J Biol Chem 279, 2101–2108 (2004).

46. K. Bobik, J. R. Dunlap, T. M. Burch-Smith, Tandem high-pressure freezing and quick freeze substitution of plant tissues for transmission electron microscopy. J Vis Exp, e51844 (2014).

47. A. W. Robards, in Intercellular Comminucation in Plants: Studies on Plasmodesmata, B. E. S. Gunning, A. W. Robards, Eds. (Springer-Verlag Berlin Heidelburg New York, Heidelburg, 1976), chap. 2, pp. 15–57.

48. T. M. Burch-Smith, P. C. Zambryski, Loss of INCREASED SIZE EXCLUSION LIMIT (ISE)1 or ISE2 increases the formation of secondary plasmodesmata. Curr Biol 20, 989–993 (2010).

49. M. Alazem, K. Y. Lin, N. S. Lin, The abscisic acid pathway has multifaceted effects on the accumulation of Bamboo mosaic virus. Mol Plant Microbe Interact 27, 177–189 (2014).

50. M. G. Johnston, C. Faulkner, A bootstrap approach is a superior statistical method for the comparison of non-normal data with differing variances. The New phytologist 230, 23–26 (2021).

