## Supplemental Material for "High-resolution microscopy of whole plant cells reveals virus-induced changes to intercellular connectivity"

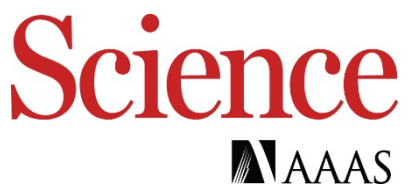

### Supplementary Materials for

#### **High-resolution microscopy of whole plant cells reveals virus-induced changes to intercellular connectivity**

Brandon C. Reagan<sup>1, †</sup>, Allyson Angermeier<sup>2</sup>, Samantha P. Nuzzi<sup>2,3</sup>, Mazen Alazem<sup>2</sup>, Andrea A. Zanini<sup>2</sup>, Alejandro Lugo Saaverda<sup>2,4</sup>, Colten Nichols<sup>2,5</sup>, Lolita Rotkina<sup>2</sup>, Kirk J. Czymmek<sup>2</sup>,
Tessa M. Burch-Smith<sup>1,2</sup>

##### **This PDF file includes:**

Materials and Methods

Figs. S1 to S11

Table S1

### Materials and Methods

#### Plant growth

*Nicotiana benthamiana* plants were grown on light carts under long day conditions (16hrs light and 8 hours of dark, 80-120 $\mu\text{mol/m/s}$). Seedlings were germinated and transferred to individual pots 10-12 days post germination. All plants used for experiments were at least 4-weeks old. Fertilizer was applied once, 5 days after transplanting.

#### Viruses and virus infection

For viruses with infectious clones (TMV, TRV, TuMV-GFP, ACMV, and EACMCV), two leaves from 4-week-old plants were agroinfiltrated with *Agrobacterium tumefaciens* containing the corresponding infectious clone. Infiltrations were done at an OD of 0.1 to ensure that most cells within the infiltrated leaf were infected with virus. All experiments with systemically infected leaves were completed 10 days post infection. Infection was confirmed via RT-PCR with primers for the coat protein for each virus (fig. S1 and table S1).

#### Sample Preparation for FIB-SEM

Tobacco samples were prepared as described previously (1) based on a modified protocol (2). Briefly, tobacco leaves (~5 mm x 5 mm) were fixed in 2% paraformaldehyde, 2% glutaraldehyde, 0.01% Tween-20, 0.05% Malachite green in 0.1M sodium cacodylate buffer (pH 7.4) at 4°C overnight. Samples were then rinsed, fixed in 1% osmium tetroxide 0.1M sodium cacodylate buffer for 4 hours followed by treatment with 2.5% potassium hexacyanoferrate(II) trihydrate in 0.1M sodium cacodylate buffer, 1% thiocarbohydrazide and subsequently 2% osmium tetroxide. Following fixation, samples were sequentially *en bloc* stained with aqueous uranyl acetate and lead aspartate before dehydrating in graded series of acetone and propylene oxide before embedment in a hard formulation Quetol 651 (Electron Microscopy Sciences).

#### FIB-SEM Imaging and Data Collection

All samples were mounted on aluminum pins (Catalog no. 16145, Thermo Fisher Scientific) rendered conductive via silver epoxy and/or metal coating (fig. S1a) and prepared and imaged on a Helios 5 Hydra CX DualBeam Plasma FIB-SEM (Thermo Fisher Scientific) with focused plasma beams for site preparation (fig. S1b). The area of interest – one epidermal cell with convoluted intercellular boundary which is not immediately next to the guard cells or the trichome cells- was selected.

The surface of the selected cell with the adjacent area was protected with Pt GIS with 12 kV Xenon for smoother surface to minimize the curtaining during long milling cycles.

After protection, the first cut was made and the trench milled (fig. S1c). The trenching was done by using Xe at 30 kV, with current varying depending on the area and the pattern size and the strategy for the entire area of interest.

When the area prep and trenching was completed, the Oxygen and set ASV (Thermo Scientific™ Auto Slice & View™ 5 (AS&V) Software) were used for automated serial section milling followed by imaging. The Multidetector setting was used to collect two sets of images simultaneously - secondary (TLD SEM) and backscattered (ICD). The milling and imaging conditions for each dataset provided herein are detailed in Table S2 and were collected using AutoSlice&View workflow (Thermo Scientific™ Auto Slice & View™ 5 (AS&V) Software). To capture the entire cell volume several partial cell volumes were collected. Image datasets were deposited in the Electron Microscopy Public Image Archive (EMPIAR) under accession number EMPIAR-13966.

#### **Image Segmentation and Rendering**

FIB-SEM images were imported in Dragonfly 3D World version 2025.1 (Comet Technologies Canada Inc., Montréal, Canada) for visualization, segmentation, and analysis. Images were manually aligned using the slice registration function. Cell walls were periodically manually segmented after which the interpolation function was used to fill segmentation gaps. PD were marked by creating a multi-slice ROI segmentation. The viroplasm and ER (fig. S3) were segmented manually. Segmented objects were rendered in 3D and used for volumetric analyses. PD X, Y, Z coordinates were measured using the center of mass function in Dragonfly and exported for further analysis. Each PD mark was converted to a multi-ROI ensuring all PD were measured as separate objects. Modeling for figure generation, all segmented ROIs were converted to meshes with 2 iterations of smoothing applied.

#### **AI-assisted code generation**

Claude (Anthropic; version Opus 4.8) was used to develop Python code for the clustering analyses of PD. The desired analyses and data input was specified by the authors. The authors reviewed the code and outputs. The code is deposited in a GitHub repository and can be accessed here through Zenodo ([10.5281/zenodo.22812106](https://doi.org/10.5281/zenodo.22812106)).

#### **Spatial analysis of plasmodesmata**

After the PD coordinates were exported from the Dragonfly software, they were projected onto a two-dimensional plane to create a wall plane in X and Y. A convex hull was applied to define the analysis region, of which PD density was calculated as PD number divided by the convex hull area. Spatial organization and clustering was addressed using several methods. The Clark-Evans index was calculated as the ratio of nearest-neighbor mean distance to expected mean distance under complete spatial randomness (CSR) to determine if PD were clustered. Ripley's K was calculated and transformed to the L-function so that PD clustering was evaluated as $L(r) - r$ where $L(r) - r > 0$ suggests PD clustering. Pair correlation functions were then used to assess PD density at specific spatial points. Monte Carlo simulations were used to assess clustering significance against the CSR null model. 95% CSR envelopes for the L-function were defined by the 2.5<sup>th</sup> and 97.5<sup>th</sup> percentiles of the simulated curves. For each wall tested, 199 random point patterns were generated. PD clustering was assessed using density-based spatial clustering of applications with noise (DBSCAN). A minimum PD cluster size of three was applied and $\epsilon = 0.5$ $\mu\text{m}$. Of the cell walls analyzed, the epidermal-epidermal PD count is 75 for uninfected and 140 for TMV-infected. The epidermal-mesophyll walls were 33 for uninfected and 32 for TMV-infected. These analyses were done in Python 3.14 using NumPy, pandas, SciPy, scikit-learn, Matplotlib, and openpyxl.

#### **PD cluster density measurements and confocal microscopy**

Images were collected with a Leica SP8 confocal microscope (Leica Microsystems, Boston, MA, USA) with White Light Laser with filters for 488nm excitation of GFP (PDLP1-GFP or MP30-GFP) and 587nm for excitation of mCherry (PDCB1-mCherry), and emission between 500-515 for GFP and 601-618nm for mCherry. PD cluster density analyses using confocal microscopy were completed as previously described (3).

##### ***Image acquisition for PD density measurements***

Images of epidermal cells of the abaxial surface of the leaf were collected 72 hours post inoculation (hpi) as previously described (3). Leaf sections were vacuum infiltrated with water and mounted on slides for imaging. 15 $\mu\text{m}$ z-stacks were collected with a step size of 1 $\mu\text{m}$. A minimum of 15 stacks were collected for each sample for three biological replicates. The steps for image processing are shown in Fig. S7. For quantification of PD cluster distributions, maximum projections were generated for each stack and the cell wall was manually traced using ImageJ (4). Cell wall area was approximated by multiplying the total length of cell wall measured by the thickness of the z-stack (15 $\mu\text{m}$). Images were converted to 8-bit images and inverted. A binary image was generated for each image using the thresholding tool in ImageJ. The particle analyzer tool in ImageJ was used to quantify the number of PD clusters in each image. PD cluster density was calculated by dividing the number of PD clusters by the cell wall area.

##### ***Primary infected leaf experiments***

For experiments in the inoculated leaf, 4-week-old *N. benthamiana* plants were co-agroinfiltrated with an agrobacterium strain carrying a fluorescently labeled PD marker (PDCB1-mCherry) for experiments with GFP-tagged viruses and (PDLP1-GFP, MP17-GFP, or MP30-GFP) for experiments with non-tagged viruses and either empty agrobacterium or an infectious clone of the virus.

##### ***PD cluster density in upper leaves before systemic virus movement***

For experiments with systemic leaves before infection, young leaves of the infected plants were agroinfiltrated with PDCB1-mCherry or PDLP1-GFP and imaged 84 hpi.

##### ***PD cluster density in systemically infected leaves***

For systemically infected leaves, upper leaves of the infected plants were agroinfiltrated with the corresponding PD markers 7 dpi and images were collected 10 dpi with the Leica SP8 confocal microscope.

##### ***PD cluster density in Remorin and movement protein overexpressing leaves***

Expanded leaves of 4- or 5-week-old *N. benthamiana* plants were co-infiltrated with REM-HA, REM-GFP, MP30-GFP, or PDLP1-GFP and PDCB1-mCherry. Images were collected 72 hpi.

##### ***PD cluster density in NbREM1-silenced plants***

Infiltrations for VIGS were performed as previously described (5). Upper leaves of silenced and non-silenced plants were agroinfiltrated with the corresponding PD markers 14 days after infiltrations to induce VIGS. Images were collected 3 days later with the Leica SP8 confocal microscope.

##### **Sample fixation and transmission electron microscopy**

Samples were prepared for TEM via high pressure freezing and quick freeze substitution as described previously (6). Leaf punches were collected from newly emerged leaves of systemically infected and uninfected plants. The leaf punches coated with yeast paste for cryoprotection and were frozen with a Wohlwend Compact 02 High Pressure Freezer (Techno Trade International). Freeze substitution was completed as previously described (6). A FS solution composed of 2% Osmium Tetroxide and 0.1% Uranyl Acetate in pure acetone was used for fixation of all samples. The yeast paste was carefully removed and samples were embedded in epoxy resin (EMBed812, Epon). Embedding was conducted over 8 days according to the following protocol: 5%, 10%, 15%, 25%, 50%, 75%, 100%, 100% under vacuum, (v/v) resin/acetone for 24hrs. Samples were transferred into molds and baked for 48 hours at 60°C. Blocks were sectioned using a Leica ultramicrotome EM UC7. 70-100-nm sections were collected and mounted on carbon-coated copper slot grids. Grids were post stained with Reynold's lead citrate and imaged with a JEOL 1400 Flash TEM (JEOL USA) operating at 120kV.

##### **Quantification of plasmodesmata frequency by TEM**

Images were collected of cell walls between adjacent epidermal cells. Cell wall lengths were measured in ImageJ (4). PD counts from more than 100 $\mu\text{m}$ of cell wall were collected for each sample. The total number of PD per cell wall were counted. PD frequency (F) was calculated using the equation described in (7); (8). Briefly, the number of PD per micron of cell wall was divided was quantified by the section thickness and the radius of the PD (we assumed an average radius of 0.025 $\mu\text{m}$).

##### **Intercellular Movement Assays**

Movement assays were performed as described in (3). Briefly, fully expanded leaves of 4- or 5-week-old *N. benthamiana* plants were agroinfiltrated with strains encoding either free GFP or free mScarlet at very low optical density ($\text{OD}_{600} = 0.0001$) to insure sporadic transformation of epidermal cells. 48 hours post infiltration, z-stacks were collected with a Leica SP8 confocal microscope (Leica with the white light laser with filters for 488nm excitation of GFP and 569nm for excitation of mScarlet and emission between 500-515nm for GFP and 609-618nm for mScarlet. Intercellular movement was quantified by counting the number of layers of movement.

For REM-HA, MP30-GFP, and PDLP1-GFP overexpression experiments, leaves were co-infiltrated with agrobacterium strains encoding the corresponding fluorescently labeled protein at $OD_{600} = 0.1$ with strains encoding free fluorescent proteins delivered at $OD_{600} = 0.0001$. Images were collected 48 hours post infiltration (hpi).

#### **Cloning and DNA constructs**

Leaves from 4-week-old *N. benthamiana* plants were collected and frozen in liquid nitrogen. RNA was extracted with Trizol following the manufacturer's protocol (Thermo Fisher Scientific, Waltham, MA). RNA was DNase treated with DNA-free DNA removal kit (Invitrogen, Carlsbad, CA). cDNA was synthesized with M-MLV reverse transcriptase (Promega, Madison, WI) using oligo-dT primers and according to the manufacturer's protocol.

##### ***Cloning of NbREM1 fragment for VIGS***

*N. benthamiana* homologs for StREM1.3 were identified using the Solgenomics BLAST tool (https://solgenomics.net/tools/blast). Two highly similar homologs were identified with BLAST (NB ID for REM1.1 is Niben101Scf02910g01040.1 and REM1.2 is Niben101Scf04804g01002.1). A silencing construct targeting both homologs was designed using the Solgenomics VIGS Tool (https://vigs.solgenomics.net/). Primers were designed to amplify the VIGS fragment and add EcoRI and XbaI restriction enzyme sites to the ends of the fragment for cloning into the multiple cloning site of the pYL156 (pTRV-RNA2) plasmid for VIGS (5). Primers used for all cloning can be found in table S2.

##### ***Cloning of NbREM1.1***

*NbREM1.1* and *NbREM1.2* were previously shown to have redundant functions [33] and therefore *NbREM1.1* was selected for overexpression experiments. The full-length coding sequence for *NbREM1.1* was identified from the Solgenomics database (https://solgenomics.net). Primers were designed to amplify the CDS of *NbREM1.1* with attB sites for Gateway cloning. The amplified fragment was cloned using standard Gateway cloning procedures. The sequence integrity of the entry clone was verified by Sanger sequencing. The *NbREM1.1* CDS was cloned into Gateway destination vectors pGWB405 (C-terminal GFP fusion) and pGWB414 (C-terminal HA tag) with the LR reaction kit. Expression clones were verified with Sanger sequencing.

##### ***Cloning of MP30-GFP and MP30-V4A-GFP***

Gateway compatible primers were designed to amplify WT MP30 and MP30(V4A) with attB sites for Gateway cloning. To generate the V4A mutation, a single nucleotide substitution (T to C) was introduced with the forward primer. The amplified fragments were cloned into the pDONR-Zeo entry vector using the BP reaction kit. Sequence integrity and the introduction of the point mutation were verified by Sanger sequencing. The CDS for WT and MP30(V4A) were cloned into the Gateway destination vector pGWB405 to generate WT and MP30(V4A)-GFP following standard cloning methods. Expression clones were verified with Sanger sequencing.

#### **Measuring REM silencing**

Quantitative PCR to measure degree of gene silencing in *NbREM* and *NbSYTA*-silenced plants was performed on cDNA from silenced plants (made as described above) using a Bio-Rad CFX96 Touch Real-Time System with SYBR Select Master Mix (ThermoFisher Scientific, Waltham, MA). An initial denaturation step of 95°C for 1 minute was used, then 40 cycles of 95°C denaturation for 5 seconds, 59°C annealing for 15 seconds, and 70°C extension and data acquisition for 10 seconds. After the initial amplification, a dissociation curve was performed. The temperature was gradually increased from 65°C to 95°C, increasing 0.5°C every 5 seconds.

#### **Western Blotting**

Leaves from 4-week-old *N. benthamiana* plants were infiltrated with *Agrobacterium tumefaciens* containing clones for expression of REM1-HA and/or MP30-GFP fusion proteins. Infiltrations were done at an OD of 0.1. infiltrated leaves were collected 4 days later and total protein extracted as described previously (9). Proteins were detected with anti-GFP (Invitrogen, catalog no. A11122 ) or anti-HA (Covance, catalog no. MMS-101-P-200; clone 16B12) primary antibodies and then with appropriate secondary antibodies.

#### **Statistics**

For GFP-PDLP1 and PDCB1-mCherry-based PD cluster distribution assays, statistical significance was determined by unpaired student's t-tests. Statistical significance was determined for each independent biological replicate and across each complete data set. Statistical significance for GFP and mScarlet movement assays was determined using both Mann-Whitney U-tests and the bootstrapping method outlined in (10). For PD frequency quantification by TEM, statistical significance was determined by Mann-Whitney U-test.

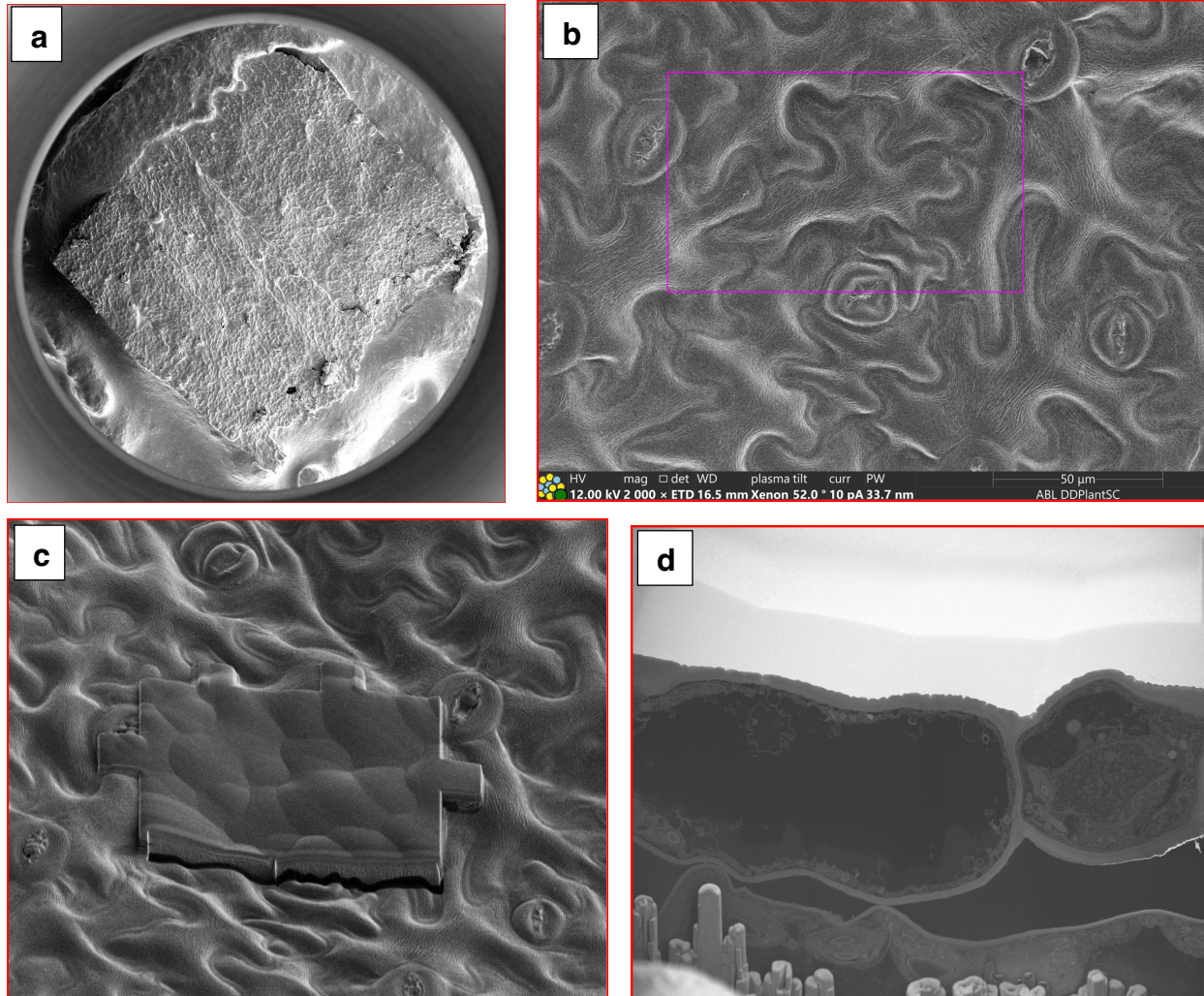

**Fig. S1. Steps for the sample preparation before automated serial sectioning.**

Portion of the sample, carved from the epoxy embedded lief, with the exposed epidermal cells mounted on the aluminum stub with the silver paste (a). The area of interest defined by using the SEM mapping over the large area and choosing the cell which is the best representation of the proposed experiment (b). The protective layer gets deposited over the area of interest (c). The connection between cells with the projected volume containing the adjacent cellular membranes (d) is chosen as a starting point for the automated serial milling.

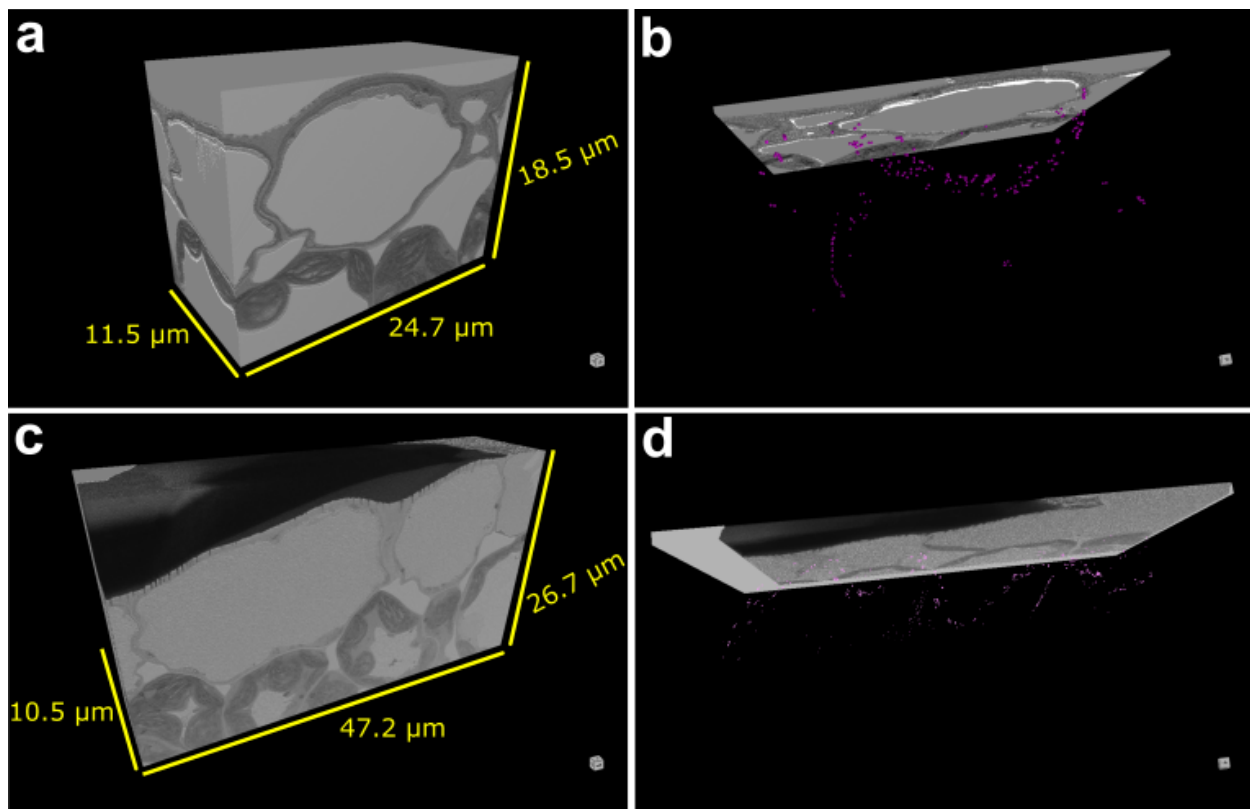

**Fig. S2. Views of whole, TMV-infected cells collected by FIB-SEM.** (A) 3D view of a *N. benthamiana* epidermal cell infected with TMV revealing general cell structures. (B) 3D view of the sample shown in (A) with the images partially scaled back to reveal PD marks (magenta dots). (C, D) 3D views of a different set of TMV-infected cells shown with complete volume shown (C) or images scaled back to reveal PD marks (Magenta dots). FIB-SEM images were collected with Adaptive Scanning Technology. The dimensions of each surface are indicated in yellow text.

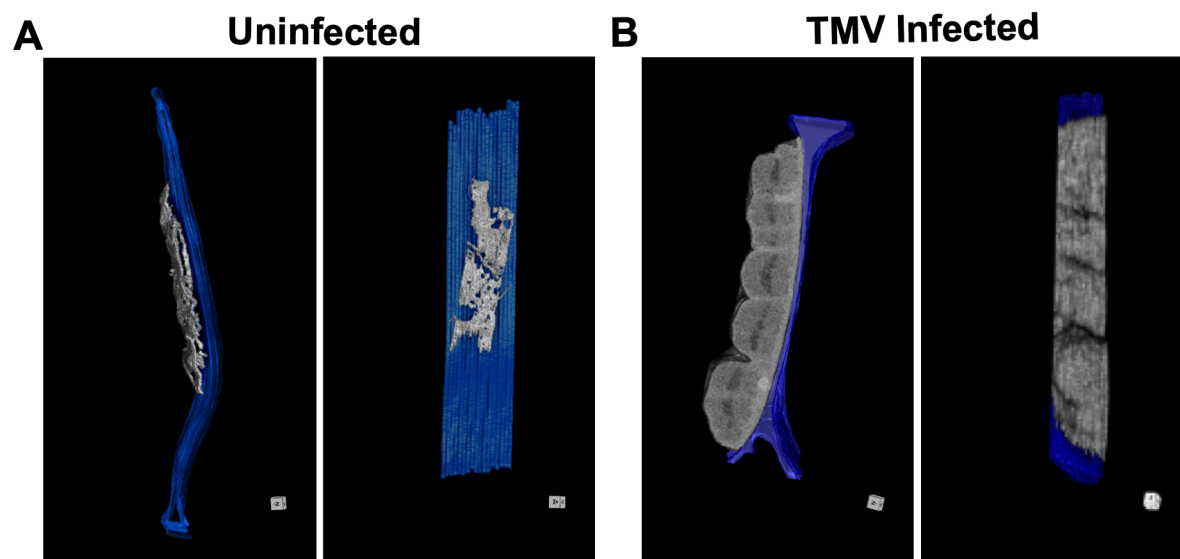

**Fig. S3. Structure of the ER in uninfected epidermal cell and the viroplasm in TMV-infected epidermal cell. (A)** 3D segmentation projection of the uninfected epidermal cell wall interface (blue) along with the ER in the Z plane (left panel) and X plane (right panel). The ER was segmented manually, applied as a mask, and used to model the ER directly from the vEM images. **(B)** 3D segmentation of the TMV-infected epidermal cell wall interface (blue) with a viroplasm. Views are the Z plane (left panel) and X plane (right panel). The viroplasm was segmented and subtracted from the vEM images as was done in (A).

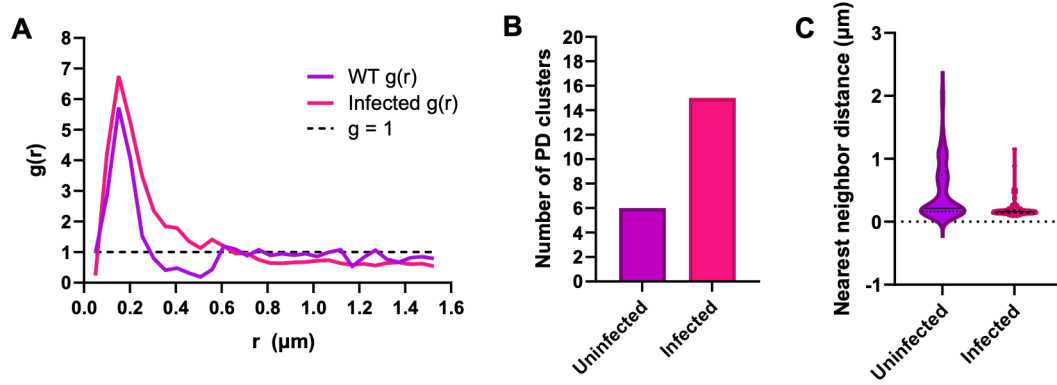

**Fig. S4. Further analysis of PD distribution in epidermal-epidermal cell walls of uninfected cells compared to TMV-infected cells.** (A) Pair correlation function $g(r)$ for PD positions on the epidermal-epidermal cell wall interface in uninfected (purple) and TMV-infected (magenta) cells. $g=1$ indicates complete spatial randomness (dashed line). (B) Comparison of the number of PD clusters in uninfected and infected epidermal-epidermal wall interfaces. (C) Nearest neighbor distance (μm) between PD in the uninfected and TMV-infected epidermal-epidermal interfaces. Violin plots show the median in the solid black line, with quartiles represented by dashed black lines.

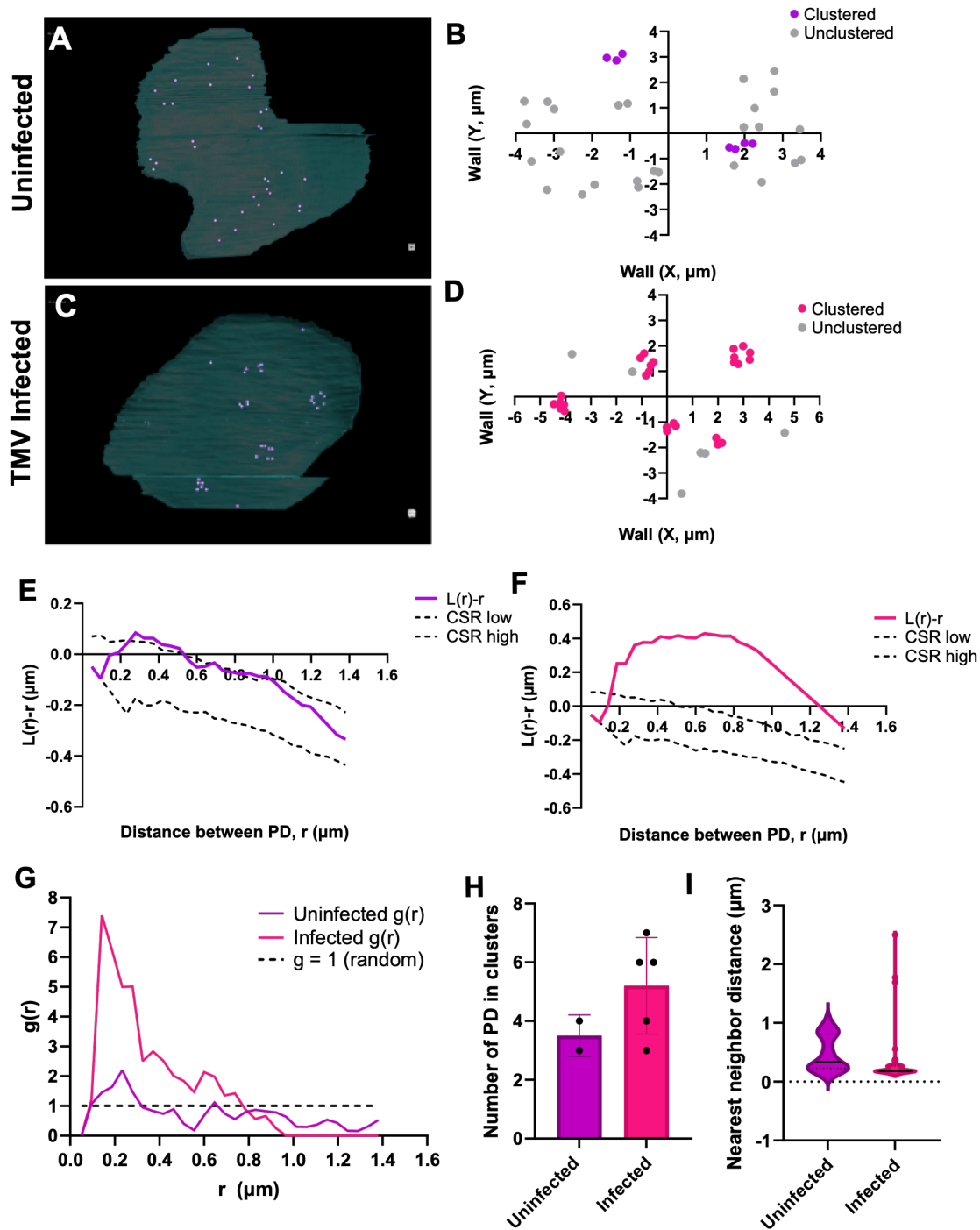

**Fig. S5. PD distribution in the uninfected and TMV-infected epidermal-mesophyll cell wall interface.** (A) Manual segmentation of the uninfected epidermal-mesophyll cell wall interface with PD (magenta dots). (B) Positions of PD on the uninfected epidermal-mesophyll interface. PD are classified as clustered (purple) or "unclustered" (grey) using DBSCAN. (C) Same as (A) for the TMV-infected epidermal-mesophyll interface. (d) Same as (b) for the TMV-infected epidermal-mesophyll interface where "clustered" PD are in magenta and "unclustered" are in

grey. (E) Spatial analysis of PD distribution on the epi-meso cell wall interface. Ripley's L-function is represented in the solid purple line, the upper and lower bounds of the CSR are determined by Monte Carlo simulations (95% confidence) and is indicated by dashed lines. (F) Same as (E) for TMV-infected epidermal-mesophyll cell wall interface, Ripley's L-function is represented by the solid magenta line. (G) Pair correlation function $g(r)$ for PD positions on the epidermal-mesophyll cell wall interface in uninfected (purple) and TMV-infected (magenta) cells. $g=1$ indicates complete spatial randomness (dashed line). (H) Comparison of the number of PD clusters in uninfected and infected epidermal-mesophyll wall interfaces, individual points indicate a cluster ($n=1$). (I) Nearest neighbor distance ($\mu\text{m}$) between PD in the uninfected and TMV-infected epidermal-mesophyll interfaces. Violin plots show the median in the solid black line, with quartiles represented by dashed black lines.

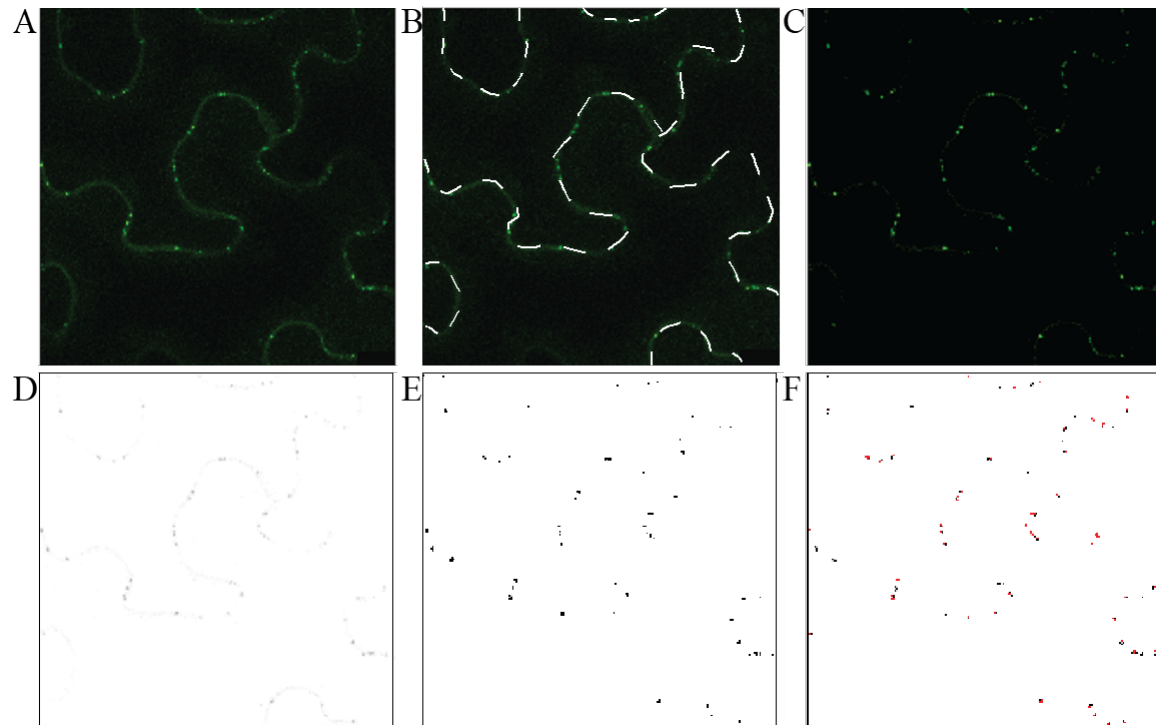

**Figure S6. Image processing pipeline for measuring PD cluster densities by confocal microscopy and fluorescently labeled PD marker proteins.** (A) A representative image of a typical maximum projection used for PD cluster density quantification is shown. (B) The total length of cell wall in the image was measured in ImageJ (dashed line). (C) Background fluorescence was decreased, and the intensity of labeled foci was maximized in ImageJ. (D) The adjusted image was converted to an 8-bit image and inverted. (E) A binary image of the inverted image was generated using the thresholding tool. (F) The particle analyzer was used to count the number of foci present in the binary image and an outline map of all counted foci was directly compared to the original image to ensure that all foci in the image were counted.

Upper leaves before infection (84 hpi)

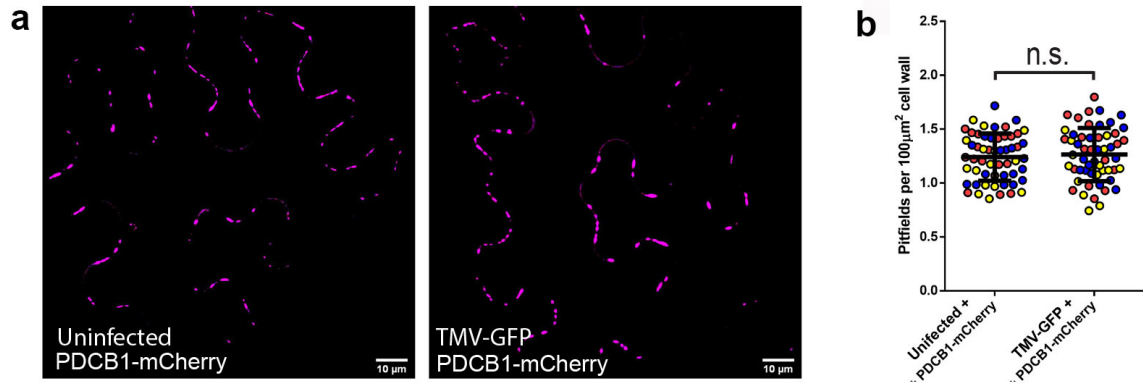

**Fig. S7. The increase in PD formation occurs only in the presence of TMV. (A)**

Representative maximum projection of z-stack of images of *N. benthamiana* upper epidermal leaf cells expressing PDCB1-mCherry with lower leaves uninfected (left) or infected TMV-GFP (right). Scale bar = 10 $\mu\text{m}$. Images were collected from upper leaves 84 hours post infection (hpi). (B) PD cluster densities were calculated using AtPDCB1-mCherry as a marker. PD cluster densities for 20 foci for each of 3 biological replicates are shown (red, blue, and yellow circles). Statistical significance was determined using Student's t-test ($p < 0.0001$).

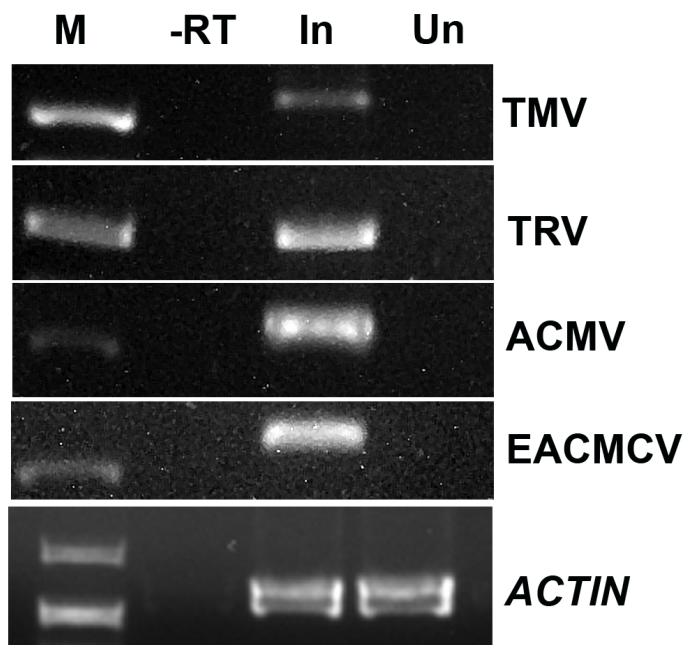

**Fig. S8. Confirmation of virus infection by reverse-transcription followed by PCR (RT-PCR).** The presence of virus in upper, systemic leaves was detected by RT-PCR with primers for the CP of TMV (First row), TRV (second row), ACMV (third row), or EACMCV (fourth row). For each virus, a single band of the expected size appears in the lane for virus infected tissue (In) and not in the negative control (-RT) or the uninfected (Un) samples. Lane M shows the band from the DNA Ladder closest in size of the targets. *ACTIN3* was used as a control.

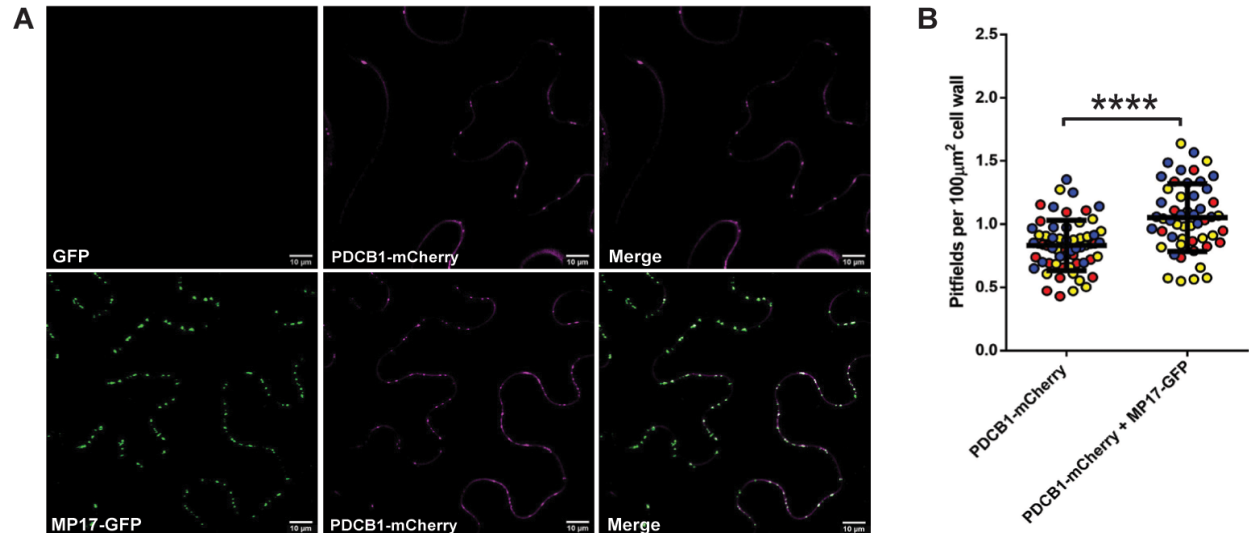

**Fig. S9. Potato leaf roll virus MP17 induces PD formation.** (A) Representative images of *N. benthamiana* epidermal leaf cells expressing either PDCB1-mCherry alone (top row) or co-expressed with MP17-GFP (bottom row). Scale bars = 10 $\mu\text{m}$. (B) PD cluster density was calculated based on the number of PDCB1-mCherry foci detected in 15 $\mu\text{m}$ z-stacks. Images were collected 96 hours post infiltration. PD cluster densities were quantified for at least 20 fields of view for 3 biological replicates (red, yellow and blue circles represent different biological replicates). Statistical significance was determined using Student's t-test ($p < 0.0001$).

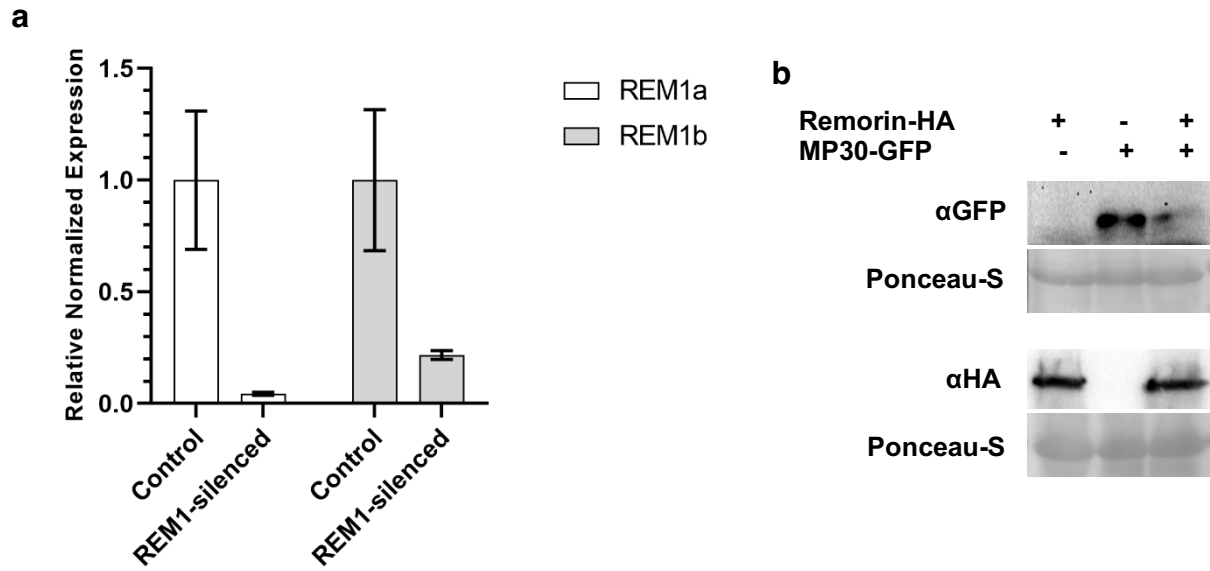

**Fig. S10. Confirmation of *NbREM1*-Silencing and expression of *NbREM1*-HA.** (A) qRT-PCR results showing that the VIGS fragment designed to target *NbREM1a* results in a 95% reduction in *NbREM1.1* expression and an 88% reduction in *NbREM1.2* expression relative to non-silencing control plants. GADPH and EF1α were used as reference genes to calculate relative expression levels normalized against the TRV-non-silencing control plants. (B) Western blot showing expression of *NbREM1*-HA. Expression was checked in tissue expressing: *NbREM1*-HA alone (Lane 1), MP30-GFP alone (Lane 2) and *NbREM1*\_HA + MP30-GFP (Lane 3). Ponceau staining of the gels is shown as the loading control.

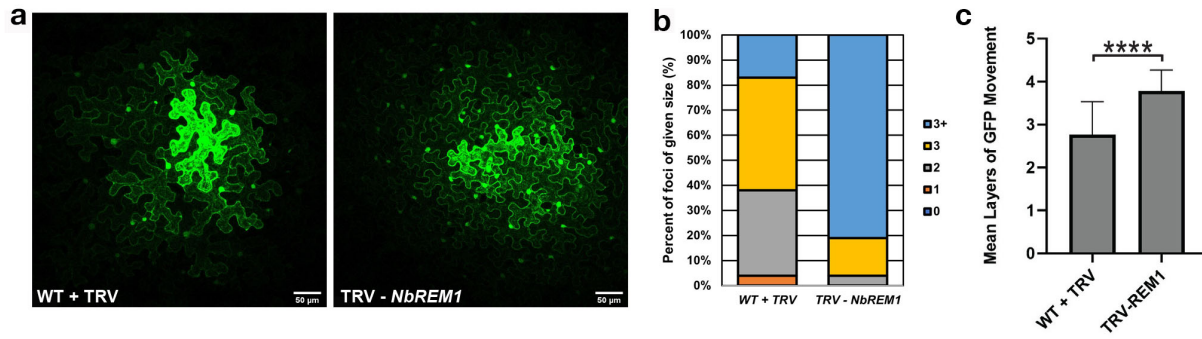

**Fig. S11. Silencing NbREM1 increases intercellular trafficking.** (A) Representative images of GFP containing foci from non-silencing controls (A, left) and *NbREM1*-silenced *N. benthamiana* leaf epidermal cells (A, right). The brightest cell at the center of the image indicates the cell transformed with the GFP marker. The GFP moves from that cell into neighboring cells producing the weaker fluorescent signals. Scale bars = 50 μm. (B, C) Intercellular trafficking was measured by counting the number of layers (rings) of cell containing GFP around a primary transformed cell. (B) Intercellular trafficking (movement) data is presented as a plot of the percentage of foci of each size. (C) Mean layers of movement and standard deviation. Data for three biological replicates are shown and at least 20 foci were collected for each biological replicate. Statistical significance was determined using the Mann-Whitney U-test ($p < 0.001$).

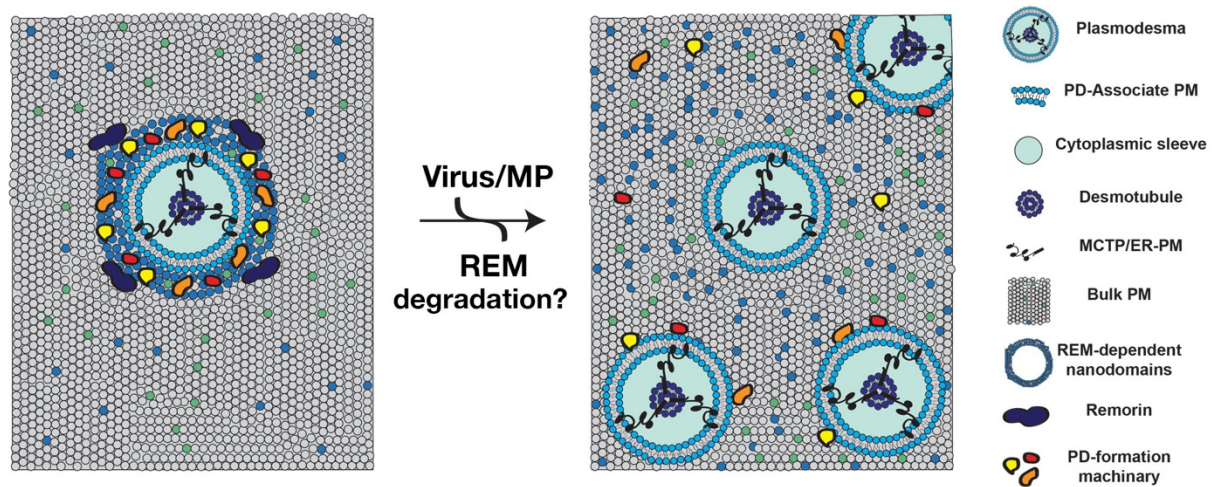

**Fig. S12. Proposed model of virus-induced PD formation.** In uninfected cells (left), REM1 organizes the PM into specialized membrane nanodomains at PD (transverse view from cytoplasm depicted). These nanodomains are enriched in long-chain fatty acids and sphingolipids and restrict *de novo* pore formation by confining membrane-associated PD formation machinery to PD membranes. During virus infection (right), REM1 is degraded, thereby disrupting PM organization. Disruption of membrane organization allows the PD-formation machinery to diffuse throughout the PM and induce unrestricted PD formation. Elevated PD formation leads to increased PD density, intercellular trafficking, and virus movement.

**Table S1. Primers used for viral detection, cloning and qPCR**

| Primer Name | Target | Primer Sequence |
| --- | --- | --- |
| TMVF | TMV CP FWD | ATGTCTTACAGTATCACTACTCCATCTCAGTTCG |
| TMVR | TMV CP RVSE | TGGGCCCCTACCGGGGGTAA |
| GeminiF | ACMV/EACMV FWD primer for CP | ATGTCGAAGCGACCAGGAGAT |
| ACMVR | ACMV RVSE primer for CP | TGTTTATTAATTGCCAATACT |
| EACMCVR | EACMV RVSE Primer for CP | CCTTTATTAATTTGTCACTGC |
| TRVF | TRV Replicase FWD | CGGAGAATGAGCTGTGGATG |
| TRVR | TRV Replicase RVSE | TGATAGAGACCTCCTCGGAAC |
| V4AF | MP30 V4A FWD attB | GGGGACAAGTTTGTACAAAAAAGCAGGCTTCATGGCTCTAGCTGT TAAAGGAAA |
| V4AR | MP30 V4A RVSE attB | GGGGACCACTTTGTACAAGAAAGCTGGGTAAAACGAATCCGATT CGG |
| MP30F | WT MP30 FWD attB | GGGGACAAGTTTGTACAAAAAAGCAGGCTTCATGGCTCTAGTTGT TAAAGGAAA |
| VIGS-REM1F | REM1 VIGS FWD EcoRI | AAAGAATTCCCATGGCAGAAGTAGAAGC |
| VIGS-REM1R | REM1 VIGS RVSE XbaI | TTTTCTAGATCTCTGTTGCAACTCGAGCAAGC |
| REM1aF | *NbREM1a* qPCR FWD | CCAACGGCTCTGTCATGATT |
| REM1aR | *NbREM1a* qPCR RVSE | CTTGCAATGGTGAGCTGTCA |
| REM1bF | *NbREM1b* qPCR FWD | TCCTGCACAGAAAGCCTCTT |
| REM1bR | *NbREM1b* qPCR RVSE | CCATCTCGCTAAGGGGATGTA |
